# *Plasmodium* ARK2-EB1 axis drives the unconventional spindle dynamics, scaffold formation and chromosome segregation of sexual transmission stages

**DOI:** 10.1101/2023.01.29.526106

**Authors:** Mohammad Zeeshan, Edward Rea, Steven Abel, Kruno Vukušić, Robert Markus, Declan Brady, Antonius Eze, Ravish Raspa, Aurelia Balestra, Andrew R. Bottrill, Mathieu Brochet, David S. Guttery, Iva M. Tolić, Anthony A. Holder, Karine G. Le Roch, Eelco C. Tromer, Rita Tewari

## Abstract

Mechanisms of cell division are remarkably diverse, suggesting the underlying molecular networks among eukaryotes differ extensively. The Aurora family of kinases orchestrates the process of chromosome segregation and cytokinesis during cell division through precise spatiotemporal regulation of their catalytic activities by distinct scaffolds. *Plasmodium* spp., the causative agents of malaria, are unicellular eukaryotes that have three divergent aurora-related kinases (ARKs) and lack most canonical scaffolds/activators. The parasite uses unconventional modes of chromosome segregation during endomitosis and meiosis in sexual transmission stages within mosquito host. This includes a rapid threefold genome replication from 1N to 8N with successive cycles of closed mitosis, spindle formation and chromosome segregation within eight minutes (termed male gametogony). Kinome studies had previously suggested likely essential functions for all three *Plasmodium* ARKs during asexual mitotic cycles; however, little is known about their location, function, or their scaffolding molecules during unconventional sexual proliferative stages. Using a combination of super-resolution microscopy, mass spectrometry, and live-cell fluorescence imaging, we set out to investigate the role of the atypical Aurora paralog ARK2 to proliferative sexual stages using rodent malaria model *Plasmodium berghei*. We find that ARK2 primarily localises to the spindle apparatus in the vicinity of kinetochores during both mitosis and meiosis. Interactomics and co-localisation studies reveal a unique ARK2 scaffold at the spindle including the microtubule plus end-binding protein EB1, lacking conserved Aurora scaffold proteins. Gene function studies indicate complementary functions of ARK2 and EB1 in driving endomitotic divisions and thereby parasite transmission. Our discovery of a novel Aurora kinase spindle scaffold underlines the emerging flexibility of molecular networks to rewire and drive unconventional mechanisms of chromosome segregation in the malaria parasite *Plasmodium*.

## Introduction

Cell division proceeds through either mitosis or meiosis, after DNA replication to enable eukaryotes to propagate, proliferate and evolve in diverse ecological niches (Drechsler and McAinsh, 2012). Cell division and chromosome segregation diverge in different eukaryotes, but both the mechanistic basis and the molecular explanation of these differences are largely unknown.

Aurora kinases (AKs) are a conserved family of spindle-associated protein kinases, with critical roles in four aspects of cell division: (I) driving mitotic/meiotic spindle assembly and disassembly, (II) regulating spindle pole structure and dynamics, (III) promoting accurate chromosome segregation, and (IV) orchestrating cellular fission at cytokinesis (Carmena et al., 2009; Willems et al., 2018) (**Fig1A**). While the Last Eukaryotic Common Ancestor (LECA) executed all these regulatory functions with a single AK, widespread gene duplication produced variable numbers of paralogues in diverse eukaryotic subgroups (Hochegger et al., 2013). Many eukaryotic lineages have retained the singular ancestral AK, including baker’s yeast, *Saccharomyces cerevisiae* (Ipl1) (Buvelot et al., 2003), the slime mould *Dictyostelium discoideum* (aurK) (Liu et al., 2008) and the intestinal parasite *Giardia intestinalis* (Davids et al., 2008; Siman-Tov et al., 2001). *Caenorhabditis spp*. (air-1 and 2) and *Drosophila spp*. (aurA and B) have two AKs (Carmena and Earnshaw, 2003). Some lineages have three AKs: for example, mammals (Aurora A to C) (Carmena and Earnshaw, 2003), flowering plants (Aurora 1 to 3)(Kawabe et al., 2005), kinetoplastid parasites (AUK 1 to 3) (Fassolari and Alonso, 2019) and apicomplexan parasites (ARK1 to 3) (Berry et al., 2018; Berry et al., 2016).

**Fig 1.**
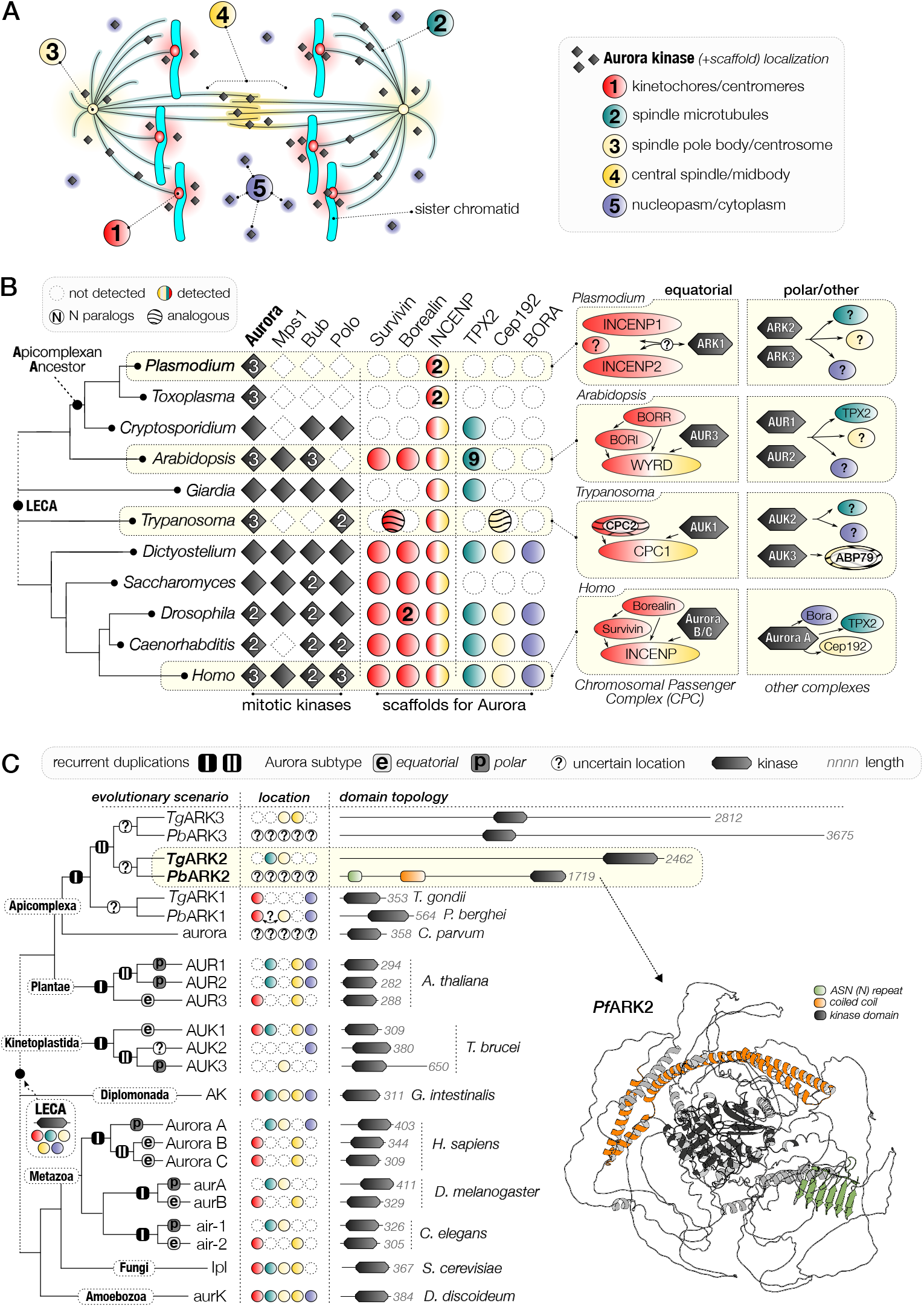
Comparative analysis of Aurora kinase family evolution reveals highly diverged paralogues amongst the Apicomplexa. **(A)** Overview of five locations of Aurora kinases (AKs) during the process of chromosome segregation in a canonical mitotic cell progression in late anaphase. Colours for each subcellular location are also used in panels B and C. **(B)** Presence-absence matrix of eukaryote mitotic kinases focused on Apicomplexa and AKs, including the scaffolds and activators of this essential kinase family. Right: (un)known mitotic location and complexes of Aurora paralogs in subgroups exemplified by model systems. LECA is last eukaryotic common ancestor **(C)** Recurrent duplication and sub-functionalisation of the AK family in model organisms throughout the eukaryotic tree of life. Left: the phylogenetic relationships of aurora paralogs. Light and dark grey boxes indicate the Aurora subtype: polar (p) or equatorial (e). I and II indicate points of recurrent duplication in the Aurora family. Middle: (un)known subcellular location of each Aurora paralog; colours correspond to those of panel A. Right: AK domain topology; note the extended length of Aurora paralogs ARK2 and ARK3 in Apicomplexa. Right bottom: AlphaFold2-predicted structure of *P. falciparum* ARK2 (https://alphafold.ebi.ac.uk/entry/O77328).

How, when and which functions are executed by each of the Aurora paralogues (either single or multiple) varies extensively between these eukaryotic lineages. Generally, a frequent division of labour between paralogues has been proposed, which appears to be correlated with the various scaffolds that direct their subcellular location and activation (Hochegger et al., 2013), and a common pattern is found in most species. One paralogue (in humans, Aurora A) is called the “polar aurora”, due to its association with centrosomal subunits including Cep192 and the microtubule-assembly factor TPX2 (Carmena and Earnshaw, 2003; Willems et al., 2018), which govern the spindle (pole)-specific location. A second paralogue (in humans, Aurora B) has been designated the “equatorial aurora” as it localises to the midplane of a dividing cell to regulate chromosome bi-orientation on the metaphase spindle to mediate cytokinesis at the last stage of cell division (Carmena et al., 2009; Hochegger et al., 2013; Willems et al., 2018). The second paralogue associates with the chromosomal passenger complex (CPC), a heterotrimeric scaffold (comprised of INCENP, Survivin and Borealin) that provides local AK activity at inner centromeres and kinetochores until metaphase, after which it translocates to microtubules of the central spindle, to orchestrate cytokinesis (Hadders and Lens, 2022; Hindriksen et al., 2017). The third paralogue provides an evolutionary platform for novelty. In humans, Aurora C is a meiosis-specific Aurora B variant (Avo Santos et al., 2011). In plants, kinetoplastids, and apicomplexans, the three AKs are less well studied, but are likely contributing to the divergent aspects of cell division in these lineages.

*Plasmodium* spp., the causative agents of malaria, belong to the phylum Apicomplexa, a group of intracellular, unicellular parasites with unusual aspects of division and multiplication. Previous phylogenetic analyses of Apicomplexa had identified three genes for Aurora Related Kinases (ARKs) 1, 2 and 3 in *Plasmodium spp*. (Reininger et al., 2011) and the coccidian *Toxoplasma gondii* (Tg) (Berry et al., 2018; Berry et al., 2016). In *Cryptosporidium spp*. only one ARK1 has been identified, suggesting an expansion of the ARK family in the common ancestor of *Plasmodium* and *Toxoplasma* (Berry et al., 2016). Broad functional characterisation of the three *Toxoplasma* ARKs identified TgARK1 as associated with the CPC component INCENP1, while TgARK2 located at centromeres (Berry et al., 2018; Berry et al., 2016). TgARK2 and TgARK3 have been shown to interact at the spindle and spindle pole, and cleavage furrow during cytokinesis, respectively. However, their associated molecular scaffolds and/or activators have not yet been characterised (Berry et al., 2018; Berry et al., 2016). Functional studies with both the human parasite *Plasmodium falciparum* and rodent parasite *Plasmodium berghei* have suggested that all three Plasmodium ARKs are likely essential for proliferation in asexual blood stage schizogony (Bushell et al., 2017; Solyakov et al., 2011; Tewari et al., 2010). Further characterisation of ARK1 and ARK3 was limited to asexual blood stages of *P. falciparum* (Berry et al., 2016), with *Pf*ARK1 shown to be potentially associated with spindle poles (Reininger et al., 2011). Nothing is known about the location or involvement of any scaffold/activator for ARK2 in *Plasmodium spp*.

Within the mosquito host, mitotic process differs substantially from that in asexual blood stage schizogony, in which closed mitosis is associated with asynchronous nuclear division that precedes cytokinesis. During male gametogenesis in the mosquito gut, rapid mitosis is characterized by three-fold genome replication from 1N to 8N. Concomitant spindle formation and chromosome segregation happens within eight minutes without nuclear division, followed by karyokinesis and cytokinesis resulting in haploid male gametes. Meiosis commences within 24 hours of fertilisation during zygote differentiation, with an initial genome duplication from 2N to 4N. Reductive divisions occur in the subsequent oocyst, through endomitotic cycles resulting in haploid sporozoites (Guttery et al., 2022; Zeeshan et al., 2020b). Our previous studies have demonstrated an unconventional toolkit of cell cycle proteins, where mitotic protein kinases and phosphatases regulate these processes (Guttery et al., 2014; Roques et al., 2015; Tewari et al., 2010). In addition to the three divergent ARKs, seven CDK-related kinases, and four divergent Nima-like kinases have been identified in the Plasmodium genome. However, *Plasmodium* spp. seem to have lost many common cell division kinases such as Bub1, Mps1 and Polo, making their complement of cell division kinases quite different from other model eukaryotes (Guttery et al., 2014; Guttery et al., 2022; Tewari et al., 2010) (**Fig1B**). In addition, the presence and role of scaffold proteins for *Plasmodium* ARKs is poorly understood.

Here, we have performed an extensive evolutionary analysis of the AK family and used *P. berghei* to identify and characterise at the functional level the plasmodial ARK2. We used fluorescent real-time live-cell imaging, antibody-based protein pulldown, bioinformatics and functional genetic studies at distinct proliferative stages within the mosquito host to reveal that ARK2 is located at the spindle and associated with a novel protein complex that includes the microtubule plus-end tracking protein, EB1. We find that both ARK2 and EB1 are critical components for spindle dynamics and the rapid cycles of chromosome segregation and are therefore crucial factors in parasite transmission.

## Results

### Evolutionary history of spindle kinases suggests divergent roles for Aurora-Related Kinase 2 and 3 in Plasmodium spp

To gain insight into the functional roles of ARKs during *P. berghei* cell division, we first re-evaluated their evolutionary history (**Fig 1A-C**). We constructed new and previously established phylogenetic profiles (Komaki et al., 2022; Kops et al., 2020; van Hooff et al., 2017) for AKs, related mitotic kinases and their location-specific scaffolds and activators in a wide variety of eukaryotes (**Fig 1B, Table S1**). We found a pervasive loss of Aurora scaffold proteins (Survivin, Borealin, TPX2, Cep192 and BORA) in the common ancestor of Plasmodium and Toxoplasma, that correlated with the loss of the widely conserved centromere/spindle kinases (Mps1, Bub1 and Polo), and the expansion of both CPC subunit INCENP (2 paralogues) and the AK family (3 paralogues). Flowering plants (*Arabidopsis thaliana*) and kinetoplastids (*Trypanosoma brucei*, Tb) have also lost to a different extent. Kinetoplastids are the only known example of organisms with the functional analogous replacement of lost subunits Survivin and Borealin by TbCPC2 (Davids et al., 2008) and the basal body scaffold TbABP67 for Cep192, respectively (Akiyoshi, 2020) (**Fig 1B, Table S1**). To explore whether patterns of AK sub-functionalization after duplication that are common in eukaryotes might also apply to ARK1 to 3, we defined five different AK subcellular locations: (I) centromere, (II) spindle microtubule, (III) spindle pole, (IV) central spindle, and (V) cyto/nucleoplasm (**Fig 1A-C**). We mapped each of the paralogues formed after the inferred duplication events found in model eukaryotes onto these locations (Hochegger et al., 2013). Our results corroborated the previously suggested pattern of recurrent sub-functionalization after the first duplication event into an ‘equatorial’ (CPC-associated) and ‘polar’ (spindle-associated) AK paralogue (**Fig 1C**). All duplications are shared between *Plasmodium* spp. and *T. gondii*, with each paralogue being one-to-one orthologous, which strongly suggests they have the similar function. TgARK1 is located at the centromere and associated with INCENP1 and 2, but it is not at the central spindle or cleavage furrow during cytokinesis, unlike TgARK3 (Berry et al., 2016). Similarly, PfARK1 is located at, or near the spindle pole (Reininger et al., 2011) suggesting that apicomplexan ARK1 is the centromere-based equatorial-like AK. The second duplication event in mammals (Aurora B: Aurora C) and plants (AUR1:AUR2) gave rise to paralogues with similar localization profiles, with the event in mammals targeting the equatorial AK, and in plants the polar/spindle AK. In kinetoplastids, this distinction is less pronounced, with only one AK (AUR1) retaining ancestral functions, and the other paralogues (AUK2/3) are highly divergent (Akiyoshi, 2020). TgARK2 and TgARK3 are both associated with the spindle or spindle pole, consistent with the pattern of sub-functionalization after duplication, although TgARK3 has an unknown function at the cleavage furrow during cytokinesis. Strikingly, both ApiARK2 and ApiARK3 are considerably larger in size (∼1500 to 3500 residues) than other AKs (∼300 to 350 residues) including ApiARK1 (**Fig 1C**). Apart from coiled-coils and asparagine-rich regions, no clear conserved sequence or structural features were identified in Plasmodium ARK2/3 indicative of binding to additional putative interaction partners (**Fig 1C**). In summary, Plasmodium ARK2 and ARK3 are highly divergent AK paralogues, but our evolutionary reconstructions strongly implicate a role for these proteins at the spindle, and/or spindle pole.

### ARK2 is expressed in the nucleus throughout the *P. berghei* life cycle

To investigate the expression and subcellular location of ARK2, we generated a transgenic parasite line by single crossover recombination at the 3’ end of the endogenous *ark2* locus to express a C-terminal GFP-tagged fusion protein **(Fig S1A)**. PCR analysis of genomic DNA using locus-specific diagnostic primers indicated correct integration of the GFP tagging construct **(Fig S1B)**. ARK2-GFP parasites completed the full life cycle, with no detectable phenotype resulting from the GFP tagging. Expression and location of ARK2-GFP were assessed by live cell imaging; ARK2-GFP was observed in all developmental stages including asexual (blood schizogony and sporogony) **(Fig S1C, D)** and sexual (gametogony and ookinete development) (**Fig S1E, F**) stages. ARK2-GFP showed a punctate nuclear pattern with one or two focal points during blood schizogony (**Fig S1C**) and sporogony (**Fig S1D**). It was present at a single focal point with an additional more diffuse nuclear location during early stages of male gametogony (30 sec after activation) and in the zygote (2h after fertilization) (**Fig S1E, F**), but in later stages it had a more dynamic location on the spindle and spindle pole as described in the next sections. Interestingly, ARK2-GFP was not detected in mature asexual (merozoites and sporozoites) and sexual (male gametes and ookinetes) stages of development (**Fig S1C-F**).

### Spatiotemporal dynamics of ARK2-GFP during male gametogony demonstrates its association with rapid spindle dynamics

ARK2 expression was analysed during the rapid mitosis of male gametogony to understand its spatiotemporal dynamics in real time. Prior to gametocyte activation, ARK2-GFP was detected as a diffuse signal within the nucleus of most gametocytes although some had a single concentrated focus (**Fig 2A**). One minute after activation, the protein was concentrated at a single point in the nucleus, before extending into an elongated bridge-like structure, which collapsed into two separate points within the next one to two minutes (**Fig 2B, Video S1**). Each separate point extended into a bridge before collapsing and resulting in four separate foci (**Fig S1G, Video S2**). A repeat of this cycle resulted in eight foci, all within 8 minutes (**Fig S1H, Video S3**). Once mature male gamete formation (exflagellation) began, these foci faded, leaving a diffuse nuclear signal (**Fig 2A**). These cycles of extension and collapse to individual foci often started and finished asynchronously with respect to other similar events of spindle bridge and foci within a single nucleus – suggesting that mitosis in male gametogony is an asynchronous form of cell division (**Fig 2A**). The events of spatiotemporal localization of ARK2-GFP were consistent with their phenotype **(Fig S2A)**.

**Fig 2.**
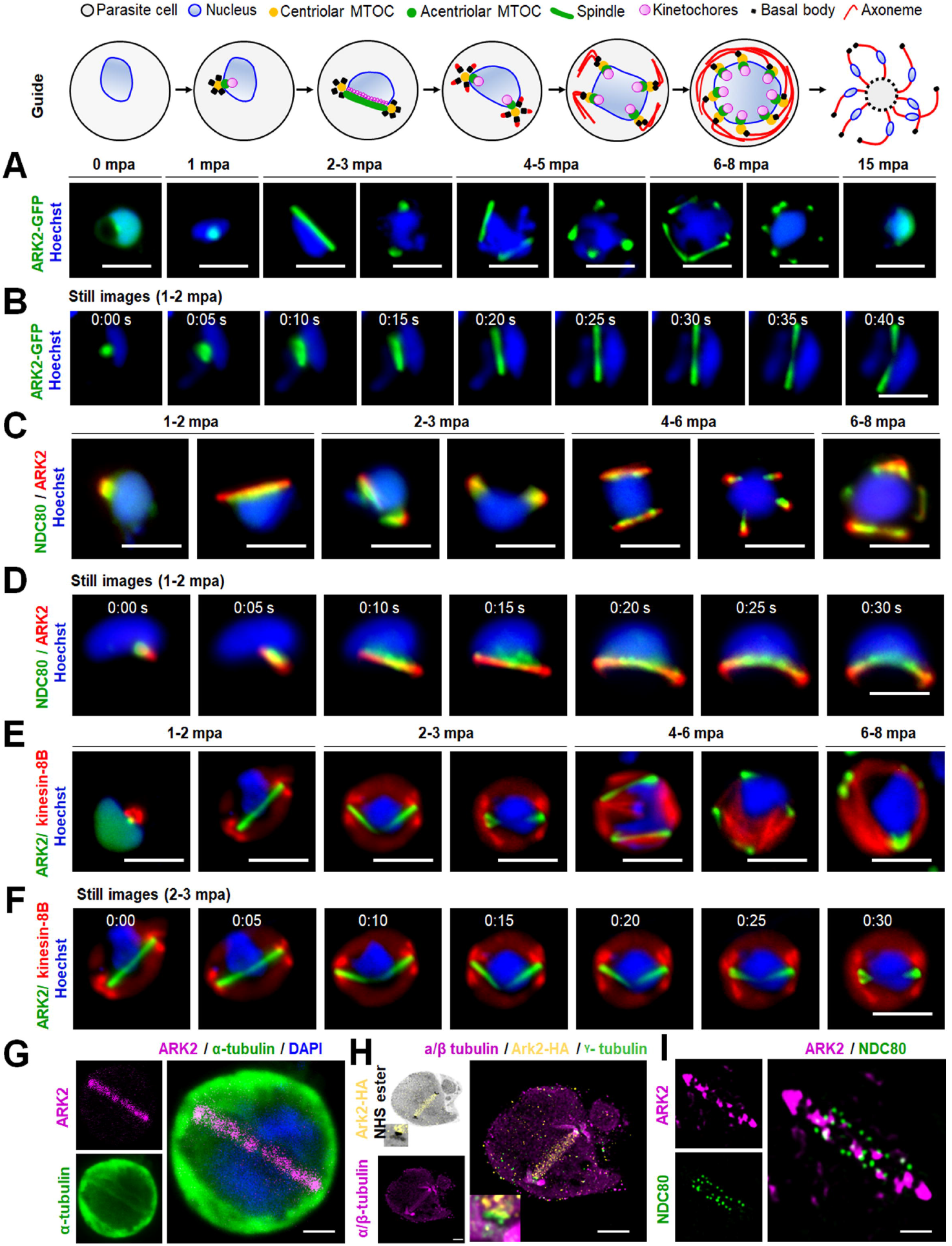
Real time live-cell imaging of PbARK2 reveals spindle association and kinetochore dynamics during male gametogony. The upper schematic shows the major stages of male gametogony with subcellular structures identified. (A) Imaging of ARK2-GFP (green) during male gametogony reveals an initial location at the putative microtubule organizing centre (MTOC) just after activation (1 minute post activation; mpa), and at the spindles and spindle poles in later stages. The protein accumulates diffusely in the residual nuclear body after gamete formation and is not present in the flagellate gametes (15 mpa). Scale bar = 5 μm. (B) Still images (5 sec intervals) showing development of an ARK2-GFP bridge from one focal point followed by further division into two halves within 1 to 2 mpa. Scale bar = 5 μm. (C) The location of ARK2-mCherry (red) relative to kinetochore marker, NDC80-GFP (green). Scale bar = 5 μm. (D) Still images (5 s intervals) of the dynamic location of ARK2-mCherry and NDC80-GFP between 1 and 2 min of activation. Scale bar = 5 μm. (E) The relative location of ARK2-GFP (green) and the basal body and axoneme marker, kinesin-8B-mCherry (red). Scale bar = 5 μm. (F) Still images (5 s intervals) of the dynamic location ARK2-GFP and kinesin-8B-mCherry between 2 and 3 min of activation. Scale bar = 5 μm. (G) Indirect immunofluorescence followed by STED confocal microscopy showing co-localization of ARK2 (purple) and α-tubulin (green) at spindle but not at cytoplasmic microtubules at 1 mpa. Scale bar = 1 μm. (H) Expansion microscopy showing co-localization of ARK2 (yellow) and α/β tubulin (purple) staining at spindle but not at cytoplasmic microtubules at 1 mpa. Scale bar = 1 μm. (I) 3D-SIM image showing locations of ARK2 (purple) and NDC80 (green) at 1 mpa. Scale bar = 1 μm. DNA (blue) is stained with Hoechst in panels A to F and with DAPI in panel G.

To examine further the location of ARK2 we investigated by indirect immunofluorescence assay (IFA) its co-localization with microtubules (MTs) that had been labelled with an α-tubulin antibody, using fixed gametocytes at different times after activation. Alpha-tubulin antibody detected both nuclear mitotic spindles and developing cytoplasmic axonemes, but ARK2 colocalized only with the mitotic spindles at all stages of gametogony (**Fig S2B**). This result provides evidence that ARK2 is involved in mitosis within the male gametocyte. To improve the resolution of detection, we used deconvolution microscopy and confirmed that ARK2 is located on mitotic spindles during male gametogony (**Fig S2C**).

### ARK2 and kinetochore dynamics are associated, but cytoplasmic axonemal microtubule dynamics are not, during male gametogony

To investigate further the association of ARK2 and the mitotic spindle during male gametogony, we compared its location with that of the kinetochore marker NDC80 and cytoplasmic axonemal protein kinesin-8B. Parasite lines expressing ARK2-mCherry and NDC80-GFP were crossed, and the progeny were analysed by live-cell imaging to establish the spatiotemporal relationship of the two tagged proteins. The location of both ARK2-mCherry and NDC80-GFP was next to the stained DNA, and with a partial overlap, although NDC80-GFP was always closer to the DNA **(Fig 2C and Fig S3A)**. This orientation of ARK2-mCherry and NDC80-GFP remained throughout male gametogony. Furthermore, the bridge length of NDC80-GFP was shorter than that of ARK2-mCherry. Time lapse imaging showed that the dynamic redistribution of ARK2-mCherry begins prior to that of NDC80-GFP and ends slightly earlier **(Fig 2D, Fig S3B, Video S4 and Video S5)**.

Parasite lines expressing ARK2-GFP and kinesin-8B-mCherry were crossed and examined by live cell imaging of both markers. One to two minutes after gametocyte activation, ARK2-GFP was observed close to the DNA and adjacent to, but not overlapping, the kinesin-8B-mCherry tetrad **(Fig 2E and Fig S3C)**. ARK2-GFP remained distributed on spindles, while there was duplication of kinesin-8B-mCherry-labelled tetrads **(Fig 2E and Fig S3C)**. In later stages of male gametogony, ARK2-GFP remained associated with spindles and spindle poles, while kinesin-8B-mCherry showed a distinct cytoplasmic axonemal location **(Fig 2E and Fig S3C)**. This location pattern was also observed in time-lapse imaging, with no colocalisation of ARK2-GFP and Kinesin-8B-mCherry **(Fig 2F, Fig S3D, Video S6 and Video S7)**. The dynamic distribution of these two proteins demonstrates that both chromosome segregation in the nucleus, tagged with ARK2, and axoneme formation in the cytoplasm, tagged with kinesin-8B begin at a very early stage of gametogony, continuing in parallel within different compartments of the male cell.

To examine further the location of ARK2 with reference to the spindle, axoneme and kinetochore at high resolution; we used super resolution confocal stimulated emission depletion (STED) microscopy, ultrastructure expansion microscopy (UExM) and 3D-structured illumination microscopy (SIM) (**Fig 2G-I**). STED images of fixed gametocytes labelled with anti-GFP, and anti-α-tubulin antibodies revealed the ARK2 distribution on nuclear spindle MTs (**Fig 2G, Fig S4**). This visualization was further improved by UExM on fixed gametocytes labelled with anti-HA antibodies (for ARK2) and anti-α/β-tubulin antibodies (for spindle and axonemes). UExM images clearly showed the ARK2 signal overlapping with spindle MTs and not with cytoplasmic axonemal MTs (**Fig 2H, Fig S4B**). These observations further indicate that ARK2 distributes on spindle MTs. Next, we used 3D-SIM on fixed gametocytes expressing both ARK2-mCherry and NDC80-GFP, which clearly showed the ARK2 bridge across the full width of the gametocyte nucleus that is associated with punctate NDC80-labelled kinetochores (**Fig 2I, Fig S4C**).

### Tracing ARK2-GFP location during the zygote to ookinete transition indicates a role at the meiotic spindle

To characterize the location of ARK2 in meiotic (i.e. zygote/ookinete) stages, ARK2-GFP dynamics were observed in developing ookinetes over a 24 h period. At various points of ookinete development, ARK2-GFP was detected as focal points like those observed during male gametogony, as well as structures radiating into the nuclear equator (**Fig 3A**). In zygotes (2h after gametocyte activation and fertilisation), ARK2-GFP was detected at one or two foci. These foci migrated away from each other over the next 8-10h through development into stage IV ookinetes, to opposite sides of the nucleus **(Fig 3A)**. During this time, the ARK2-GFP signal appeared to radiate into the centre of the nucleus, typical of a classic metaphase spindle arrangement **(Fig 3A)**. These two foci then divided again to form four foci, before the signal faded into a diffuse distribution within nuclei of mature ookinetes **(Fig 3A)**. The location of ARK2 relative to that of the kinetochore marker, NDC80, was examined during ookinete development in parasite lines expressing ARK2-mCherry and NDC80-GFP. ARK2-mCherry was located on spindles radiating from the poles and NDC80-GFP was detected along the metaphase plate (**Fig 3B**) during stages I to III. By stage IV both ARK2 and NDC80 had accumulated at spindle poles (**Fig 3B**).

**Fig 3.**
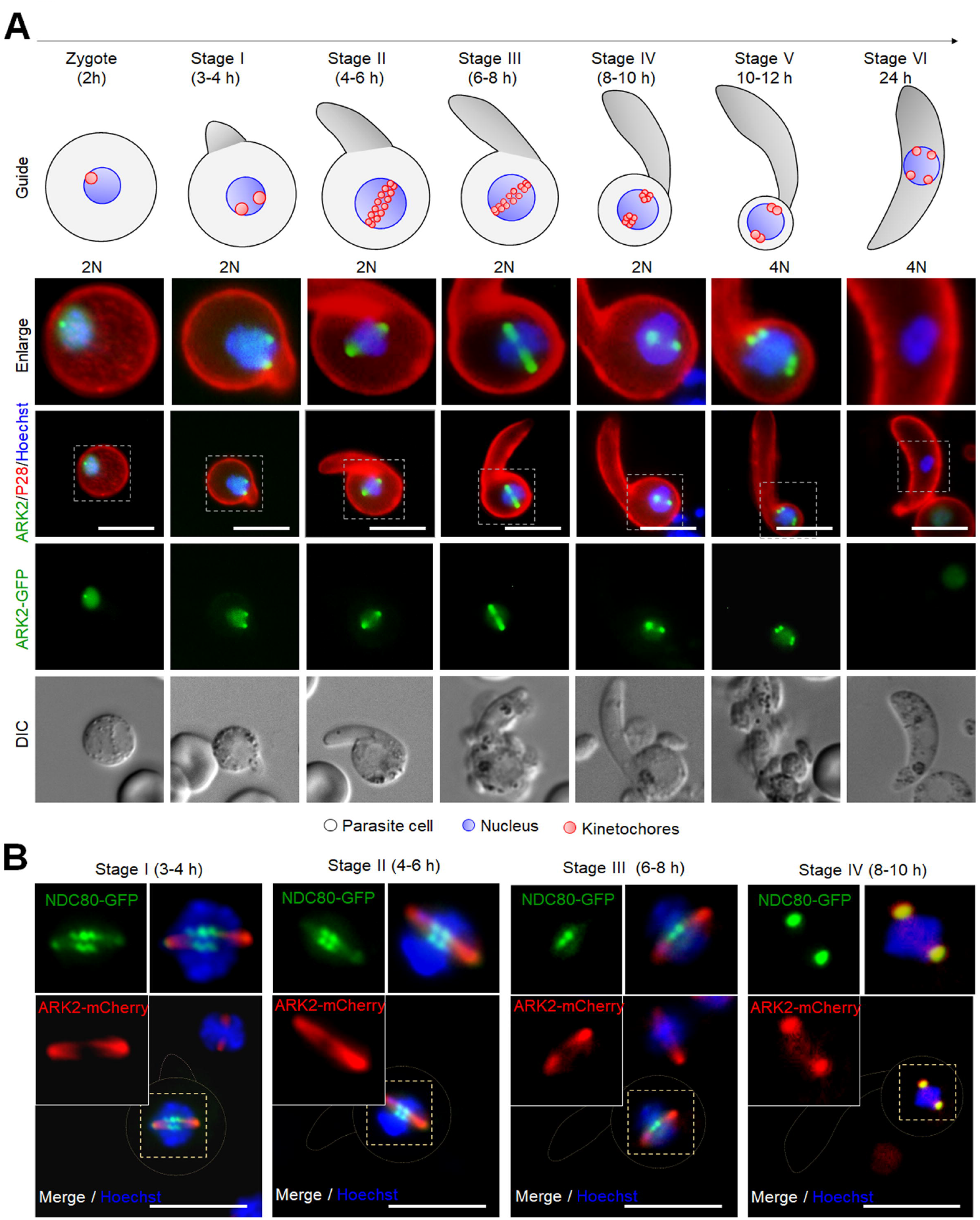
*Pb*ARK2 localizes to a putative MTOC and spindle during ookinete development. The schematic depicts ookinete differentiation from the zygote through six stages over a 24-hour period. The genome is initially diploid (2N) and then replicated (4N) just before the nucleus migrates into the growing apical protuberance. **(A)** Live-cell imaging showing ARK2-GFP (green) location during ookinete development, relative to the nuclear DNA (blue, Hoechst), and cy3-conjugated 13.1 antibody (red), which recognises P28 protein on the surface of zygotes and ookinetes. DIC images are shown in the bottom set of panels. Scale bar = 5 μm. **(B)** The location of ARK2–cherry (red) in relation to the kinetochore marker, Ndc80-GFP (green) and the nuclear DNA (blue) at different stages of ookinete development. Scale bar = 5 μm.

### Conditional knockdown of ARK2 reveals a crucial role during parasite transmission

ARK2 had previously been found to be most likely essential for asexual blood stage development (Tewari et al., 2010). To examine the role of ARK2 during sexual stages we first tagged the endogenous ARK2 locus with sequence encoding an auxin-inducible degron (AID) and an HA epitope tag (**Fig S5A**) to degrade the fusion protein in the presence of auxin in a parasite line expressing the TIR1 protein (Philip and Waters, 2015). Although the genetic modification was confirmed by diagnostic PCR (**Fig S5B**), addition of auxin to gametocytes did not lead to ARK2-AID/HA degradation **(Fig S5C**) and there was no detectable phenotype in male gametogony (**Fig S5D**). Since the AID system was unsuccessful, we used a promoter trap strategy, replacing the *ark2* promoter with that of cytoadherence-linked asexual protein (CLAG – PBANKA_1400600), which is not transcribed in gametocytes (Sebastian et al., 2012) (**Fig S5E**). The correct genetic integration was confirmed by PCR **(Fig S5F)**, and ARK2 transcription was downregulated in *P*_*clag-*_*ark2* gametocytes as shown by qRT-PCR **(Fig S5G)**. A phenotypic analysis of these *ark2*-knockdown parasites was then performed at different stages of parasite development within the mosquito.

Despite the significant reduction of ARK2 expression in *P*_*clag-*_*ark2* gametocytes (**Fig S5G**), neither mitosis in male gamete formation (exflagellation) nor meiosis in zygote differentiation (ookinete development) were affected (**Fig 4A, B**). However, serious defects in oocyst formation (endomitosis) were observed, with a significant reduction (up to 70 %) in the number of oocysts per mosquito midgut, detectable from as early as day 7 post-infection, and remaining significantly lower through to day 21 (**Fig 4C**). Microscopic imaging of the midguts revealed that the few oocysts present were smaller than those of wild-type parasites expressing GFP (WT-GFP) after day 7. Sporogony had been completely blocked; some parasites contained dark granules, and some had a pycnotic appearance (**Fig 4D**). *P*_*clag-*_*ark2* oocysts were significantly smaller than wild-type from day 14 onwards, not growing beyond the size observed at day 7 (**Fig 4E**). There were no sporozoites in the salivary glands of *P*_*clag-*_*ark2* parasite-infected mosquitoes, indicating that sporozoite development had been completely blocked even though some oocysts had formed (**Fig 4F**).

**Fig 4.**
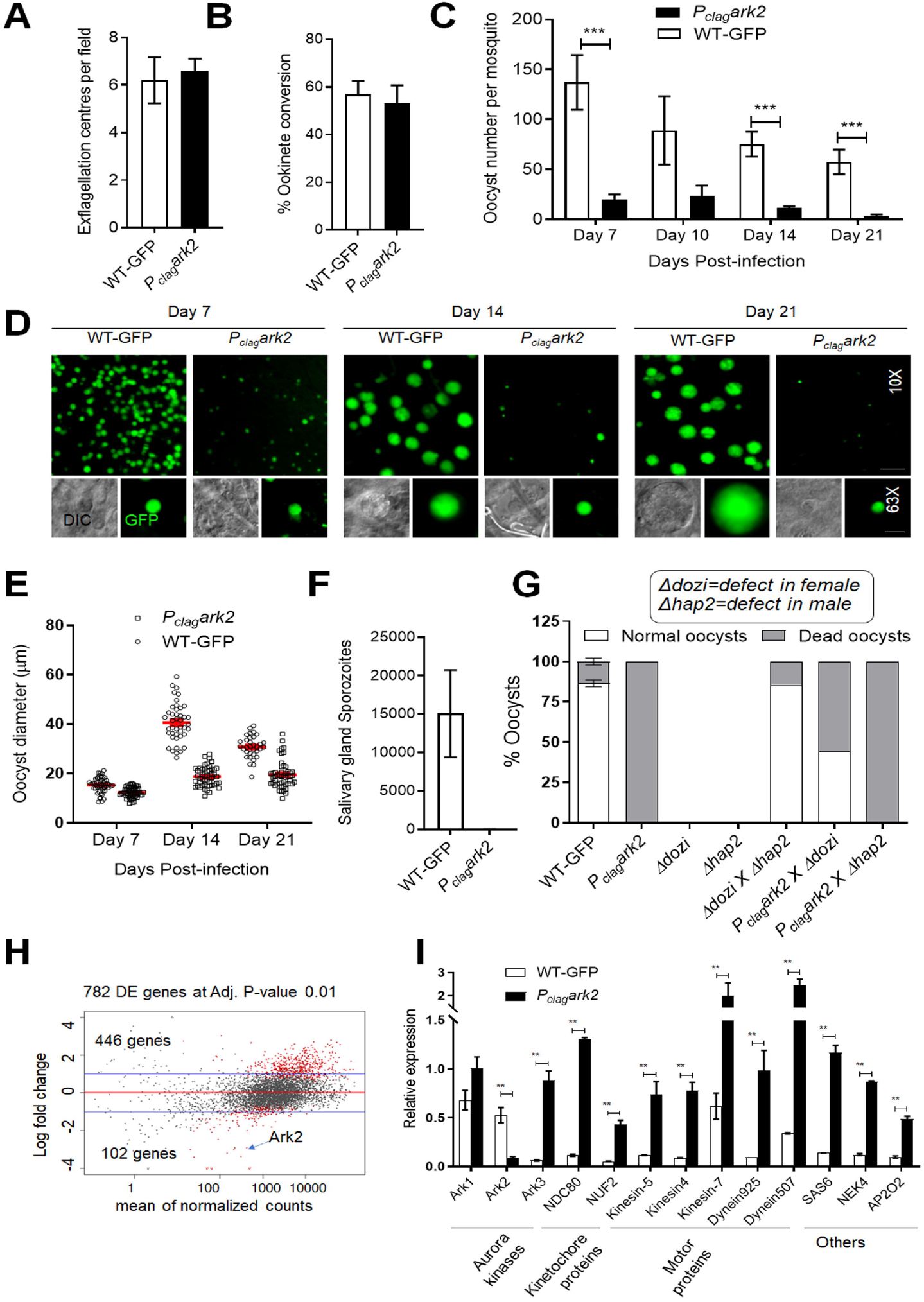
Conditional knockdown of PbARK2 identifies an essential role in oocyst development and sporogony. (A) The number of exflagellation centres per field of P_clag-_ark2 (black bar) compared with WT-GFP (white bar) parasites at the end of male gametogony. Shown is mean ± SD; n = 3 independent experiments. (B) Percentage ookinete conversion for P_clag-_ark2 (black bar) and WT-GFP (white bar) parasites. Ookinetes were identified by reactivity with 13.1 antibody and successful differentiation into elongated ‘banana shaped’ ookinetes. Shown is mean ± SD; n = 3 independent experiments. (C) Total number of GFP-positive oocysts per infected mosquito in P_clag-_ark2 (black bar) and WT-GFP (white bar) parasites at 7, 14 and 21-day post-infection (dpi). Shown is mean ± SD; n = 3 independent experiments (with >15 mosquitoes for each) ***p<0.001. (D) Mid guts at 10x- and 63x-magnification showing fluorescent oocysts of P_clag-_ark2 and WT-GFP lines at 7, 14 and 21 dpi. Scale bar = 50 μm (10x) or 20 μm (63x). (E) Oocyst sizes of P_clag-_ark2 and WT-GFP lines at 7, 14 and 21 dpi. (F) Total sporozoite number in salivary glands of P_clag-_ark2 (black bar, not visible) and WT-GFP (white bar) parasites, showing mean ± SD; n = 3 independent experiments (G) Rescue experiment showing male-derived allele of P_clag-_ark2 is affected and is complemented by ‘female’ Δdozi. (H) RNA-seq analysis showing upregulated and downregulated genes in P_clag-_ark2 parasites compared to WT-GFP parasites (I) Expression level validation of relevant selected genes from the RNAseq data using qRT-PCR. Shown is mean ± SD; n = 3 independent experiments.

One explanation for the significantly reduced number of *P*_*clag-*_*ark2* compared to WT-GFP oocysts, was reduced ookinete motility. However, when we analysed ookinete motility on Matrigel, we saw no remarkable difference in the gliding motility of *P*_*clag-*_ *ark2* **(Video S8)** compared with WTGFP parasites **(Video S9) (Fig S6A, B)**.

Since ARK2 is expressed in male gametocytes and parasite development is affected after fertilization, we investigated whether the defect is due to inheritance from the male gamete. We performed genetic crosses between *P*_*clag-*_*ark2* parasites and other mutants deficient in production of either male (Δ*hap2)* (Liu et al., 2008) or female (Δ*dozi)* gametocytes (Mair et al., 2006). Crosses between *P*_*clag-*_*ark2 and* Δ*dozi* mutants produced some normal-sized oocysts that were able to sporulate, showing a partial rescue of the *P*_*clag-*_*ark2* phenotype (**Fig 4G**). In contrast, crosses between *P*_*clag-*_*ark2 and* Δ*hap2* did not rescue the *P*_*clag-*_*ark2* phenotype. These results reveal that a functional *ark2* gene copy from a male gamete is required for subsequent oocyst development.

### Transcriptome analysis of P_clag-_ark2 parasites reveals altered expression of genes for proteins involved in several functions including microtubule-based motor activity

To explore the effect of ARK2 knockdown on the expression of other genes in gametocytes, we performed RNA-seq transcriptomic analysis of *P*_*clag-*_*ark2* and wild-type cells immediately prior to gametocyte activation (0 min) and after exflagellation (30 min post activation). The genome-wide read coverages for the four pairs of biological replicates (WT, 0 min; WT, 30 min; *P*_*clag-*_*ark2*, 0 min; and *P*_*clag-*_*ark2*, 30 min) exhibited Spearman correlation coefficients of 0.961, 0.939, 0.972 and 0.930; respectively, validating the reproducibility of the experiment. The downregulation of *ark2* gene expression in *P*_*clag-*_*ark2* gametocytes was confirmed by the RNA-seq analysis: the number of reads mapped to this gene was significantly decreased (**Fig S6C**).

In addition to changed ARK2 expression, we detected 446 and 102 genes that were significantly upregulated and downregulated respectively in *P*_*clag-*_*ark2* gametocytes activated for 30 min (**Fig 4B and Table S2**). Gene ontology (GO) enrichment analysis of the upregulated genes identified genes involved in microtubule-based processes—including microtubule-dependent motors—together with other functions including cell division and chromosome organization **(Fig S6D)**. These differences in transcript levels revealed by RNA-seq analysis were validated by qRT-PCR, focusing on genes for proteins involved in motor activity, other AKs, kinetochore proteins and genes for proteins implicated in ookinete and oocyst development **(Fig 4I)**. The modulation of these genes suggests the involvement of ARK2 in mitosis in male gametocytes, although the effect manifested only later during sporogony.

### ARK2 interacts with microtubule-binding proteins near the spindle-kinetochore interface

Until now, no scaffold or activator proteins that associate with apicomplexan ARK2 orthologues have been described. We therefore aimed to identify candidates interacting with ARK2. We first performed an immunoprecipitation experiment using anti-GFP trap beads on extracts of gametocytes expressing ARK2-GFP or GFP alone and activated for 1 min (when the first spindle is formed as described above). Lysates were prepared in the presence of limited amounts of cross-linking paraformaldehyde to stabilise protein complexes (**Fig 5A**). Immunoprecipitated proteins were then digested with trypsin prior to identification by mass spectrometry. Comparative proteomic analysis, using principal component analysis (PCA) of the GFP control and ARK2-GFP precipitates, revealed that ARK2 associates with several microtubule-associated proteins located at, or near the spindle and kinetochore (**Fig 5B, Table S3**). Generally, few unique peptides were identified for each protein except for ARK2 itself, suggesting that their binding to ARK2 may be transient or that much of ARK2 is not bound to other proteins at this stage. The interacting proteins identified include the spindle MT-associated proteins kinesin-8X (PBANKA_0805900), myosin K (PBANKA_0908500), the MT plus-end tracker EB1 (PBANKA_0405600), a variety of kinetochore proteins such as members of the NDC80 complex (Zeeshan et al., 2020b) and the recently discovered highly divergent Apicomplexan Kinetochore proteins (AKiT) AKiT1-6, STU2 (PBANKA_1337500), Mad1 (PBANKA_0612300) (Brusini et al., 2022), and a single peptide for the CPC subunit INCENP2 (PBANKA_1343200) (**Fig 5B, Table S3**). Interestingly, the strongest evidence for an ARK2 interaction was obtained for the nuclear formin-like protein MISFIT, a key regulator of ookinete-oocyst transition (Bushell et al., 2009), and a putative regulator of actin filament dynamics. We also found peptides from subunits of the Origin of Replication Complex (ORC; e.g. ORC-1/-2/-5, Cdc6 and Cdt1), and the alpha subunits of delta and epsilon DNA polymerases, which are common contaminants of immunoprecipitates from male gametocytes, possibly due to their high concentrations in the rapid cycles of replication (**Fig 5B**).

**Fig 5.**
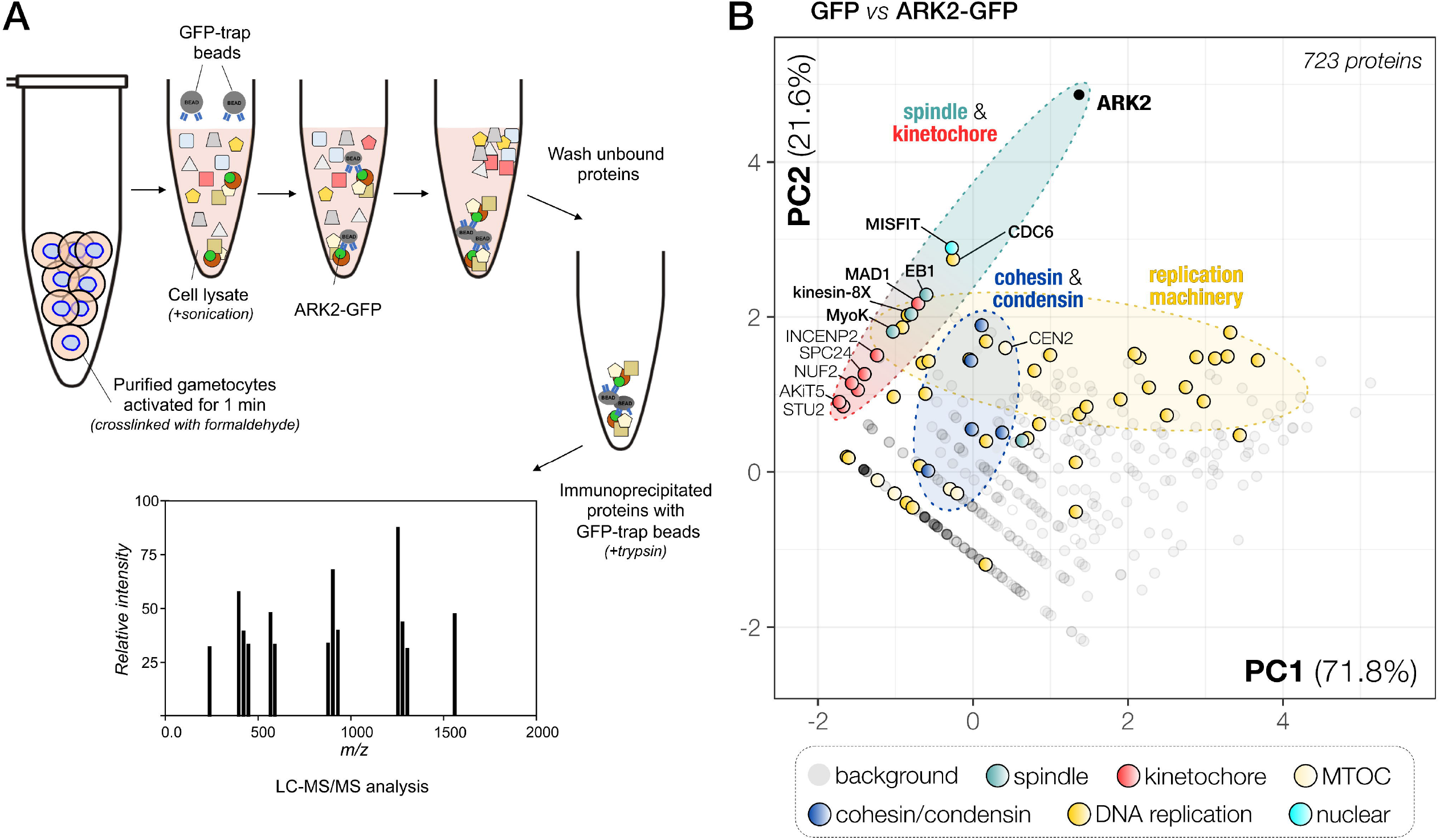
PbARK2-GFP interactome during male gametogony. (A) Workflow for immunoprecipitation experiment using GFP-trap beads and gametocyte crosslinked lysates, trypsin digestion and mass spectrometry analysis to identify ARK2-GFP interacting partners. (B) Projection of the first two components of a principal component analysis (PCA) of unique peptides derived from ARK2-GFP or GFP-alone immunoprecipitations with GFP-trap (raw data: Table S2). A subset of proteins is highlighted on the map based on relevant functional categories.

### Real time live cell imaging of parasites expressing EB1-GFP reveals its association with the spindle and kinetochore throughout male gamete formation

Our results suggested that ARK2 is located on the spindle and interacts with kinetochore components, as well as the spindle-based MT end-binding protein EB1. Therefore, we tagged EB1, encoded by the endogenous locus, with GFP (**Fig S7A, B**) and studied its spatiotemporal dynamics during male gametogony using real time live-cell imaging. EB1-GFP showed a similar spatiotemporal distribution to that of ARK2-GFP, with distinct foci and elongated spindle ‘bridges’ at certain time points after gametocyte activation (**Fig 6A, Fig S7C-E, Video S10-12**). Chromatin immunoprecipitation with parallel sequencing (ChIP-Seq) was used to determine the DNA binding sites of EB1, and indicated its co-location with the outer kinetochore marker NDC80 centromeric chromatin marker (Iwanaga et al., 2012; Zeeshan et al., 2020b) **(Fig 6B)**. These results were corroborated by live-cell imaging of EB1-GFP/NDC80-mCherry dual reporter lines **(Fig 6C, Fig S7F)**. An additional cross to produce EB1-GFP/ARK2-mCherry parasites showed overlap of fluorescence signals at 1 to 2 min post-activation of gametocytes, confirming their co-location and interaction with spindles and the kinetochore (**Fig 6D)**. Finally, parasite lines expressing EB1-mCherry and the basal body marker SAS4-GFP (Zeeshan et al., 2022) showed EB1’s association with the formation of basal bodies that serves as the MT organising centre for axonemes (**Fig 6E**).

**Fig 6.**
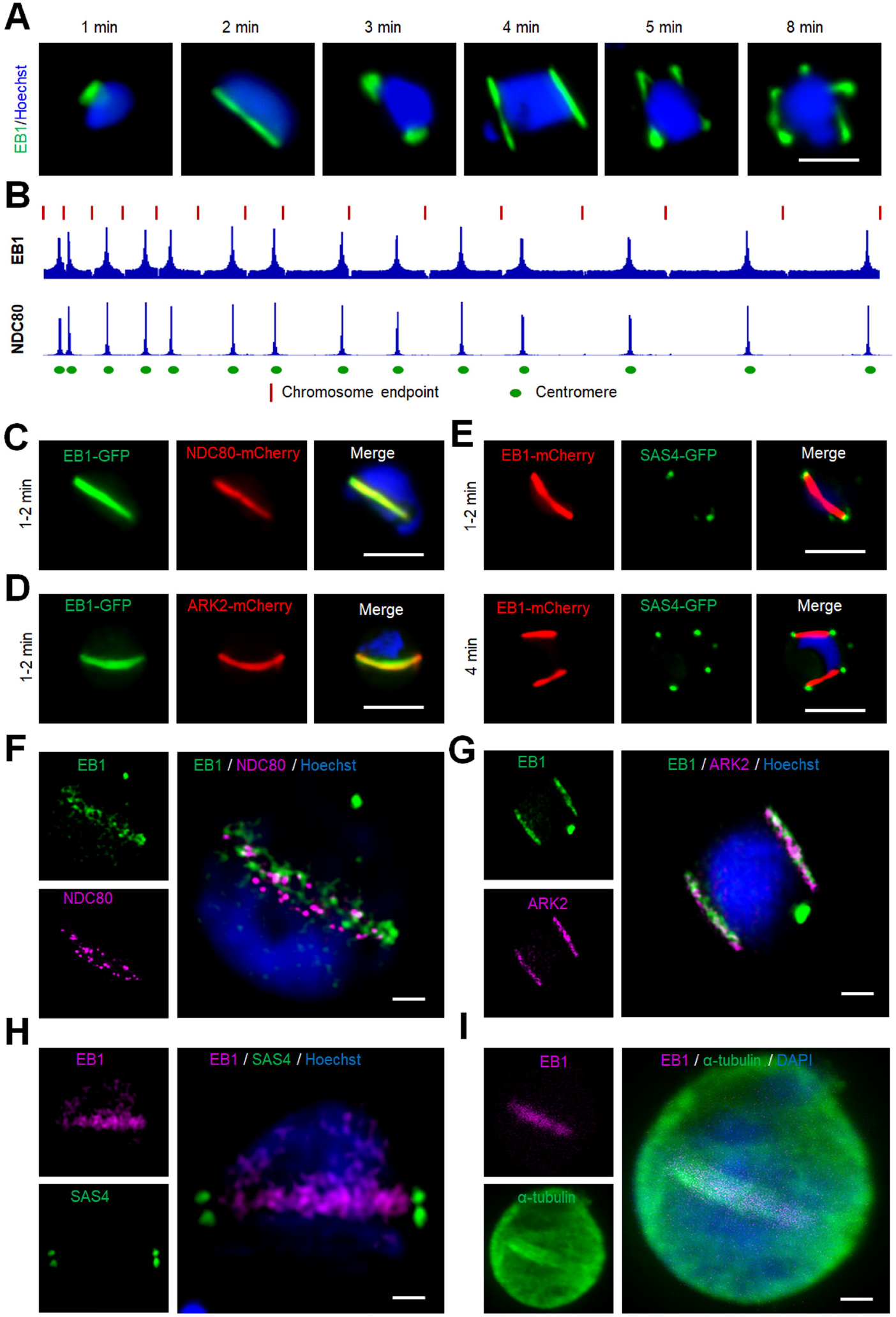
EB1 like ARK2 associates with spindle and kinetochore during male gametogony. (A) Live cell imaging of EB1-GFP (green) showing its location on spindles and spindle poles. DNA is stained with Hoechst dye (blue); scale bar = 5 μm. (B) ChIP-seq analysis of EB1-GFP profiles for all 14 chromosomes showing its centromeric binding. Signals are plotted on a normalized read per million (RPM) basis. Red lines at the top indicate the ends of chromosomes; circles on the bottom indicate centromere locations. NDC80-GFP was used as a positive control and IgG was used as a negative control. (C) Live cell imaging showing the location of EB1-GFP (green) and kinetochore marker NDC80-mCherry (red) in a gametocyte activated for 1-2 min. DNA is stained with Hoechst dye (blue); scale bar = 5 µm. (D) Live cell imaging showing the location of EB1-GFP (green) and ARK2-mCherry (red) in a gametocyte activated for 1-2 min. DNA is stained with Hoechst dye (blue); scale bar = 5 µm. (E) Live cell imaging showing the location of EB1-mCherry (red) and basal body marker SAS4-GFP (green) in gametocytes activated for 1 to 2 min (upper panel) and 4 min (lower panel). DNA is stained with Hoechst dye (blue); scale bar = 5 µm. (F) 3D-SIM image showing location of EB1 (green) and NDC80 (purple) in gametocyte activated for 1 min. DNA is stained with Hoechst dye (blue); scale bar = 1 μm. (G) 3D-SIM image showing location of EB1 (green) and ARK2 (purple) in gametocyte activated for 3 to 4 min. DNA is stained with Hoechst dye (blue); scale bar = 1 μm. (H) 3D-SIM image showing location of EB1 (purple) and cytoplasmic SAS4 (green) in gametocyte activated for 1 min. DNA is stained with Hoechst dye (blue); scale bar = 1 μm. (I) STED confocal microscopy showing co-localization of EB1 (purple) and α-tubulin (green) at spindle but not with cytoplasmic microtubules in gametocytes activated for 1 min. DNA is stained with SiR DNA (blue); scale bar = 1 μm.

To further resolve the location of EB1 with respect to the kinetochore and basal body at higher resolution, 3D-SIM was performed on EB1-GFP/NDC80-mCherry, EB1-GFP/ARK2-mCherry and EB1-mCherry/SAS4-GFP fixed gametocytes. The 3D-SIM images of gametocytes expressing EB1-GFP/NDC80-mCherry showed EB1 bridge(s) across the nucleus with NDC80 distributed like beads on the bridge, each bead representing a kinetochore (**Fig 6F, Fig S8A**). The 3D-SIM images of gametocytes expressing EB1-GFP/ARK2-mCherry showed EB1 bridge(s) across the nucleus with ARK2, overlapping each other ((**Fig 6G, Fig S8A**). The bridged pattern of spindles for EB1 were restricted to the nucleus as shown by 3D-SIM images of gametocytes expressing EB1-mCherry/SAS4-GFP; whereby SAS4 was located in the cytoplasm but aligned with the EB1 bridge in the nucleus **(Fig 6H, Fig S8A)** (Zeeshan et al., 2022). We also performed STED microscopy on fixed EB1-GFP gametocytes stained with anti-GFP and anti-tubulin antibodies, which confirmed EB1’s location on the spindle: the images showed EB1 distribution on the nuclear spindle MTs with a distribution like that of ARK2 (**Fig 6I, Fig S8B**). These real time imaging data, together with the interactome data, confirm that ARK2 and EB1 form a functional/structural axis associated with the spindle and the acentriolar MTOC, and are associated with kinetochore dynamics.

### EB1-GFP is enriched at spindles and associated with apical polarity during ookinete differentiation

Live cell imaging of EB1-GFP during early meiosis located the protein on spindles and spindle poles, but it then disappeared as the ookinete matured, with a pattern similar to that observed for ARK2-GFP (**Fig S9**). There was also an accumulation of EB1-GFP at the nascent apical end of the developing ookinete, potentially important in defining its polarity. In later stages of ookinete differentiation, it was distributed around the periphery of the growing protuberance, potentially associated with sub-pellicular MTs (**Fig S9)** but had disappeared in mature ookinetes.

### EB1 is not essential for asexual blood stage proliferation, but like ARK2 its deletion affects endomitosis during sporogony

We showed that ARK2 and EB1 have similar spatiotemporal dynamics but wanted to establish whether deletion of the EB1 gene would have a similar phenotype to that of an ARK2 mutant line. We therefore generated an EB1 gene deletion mutant (Δ*eb1*) via double homologous recombination (**Fig S10A, B**). This deletion had no effect on asexual blood stage parasite development, in contrast to ARK2 that is essential during blood schizogony (Tewari et al., 2010). Male gamete formation (exflagellation), fertilisation and zygote differentiation (ookinete development) were also not affected in Δ*eb1*parasites (**Fig 7A, B)**. However, deletion of *eb1* resulted in significantly reduced oocyst numbers on day-10 post-infection of mosquitoes (**Fig 7C)**. The oocysts that were present were smaller than those of WT parasites (**Fig 7D**), and by day-21 no oocysts were detectable (**Fig 7C, D**) suggesting that development was completely blocked at some point beyond day-10.

**Fig 7.**
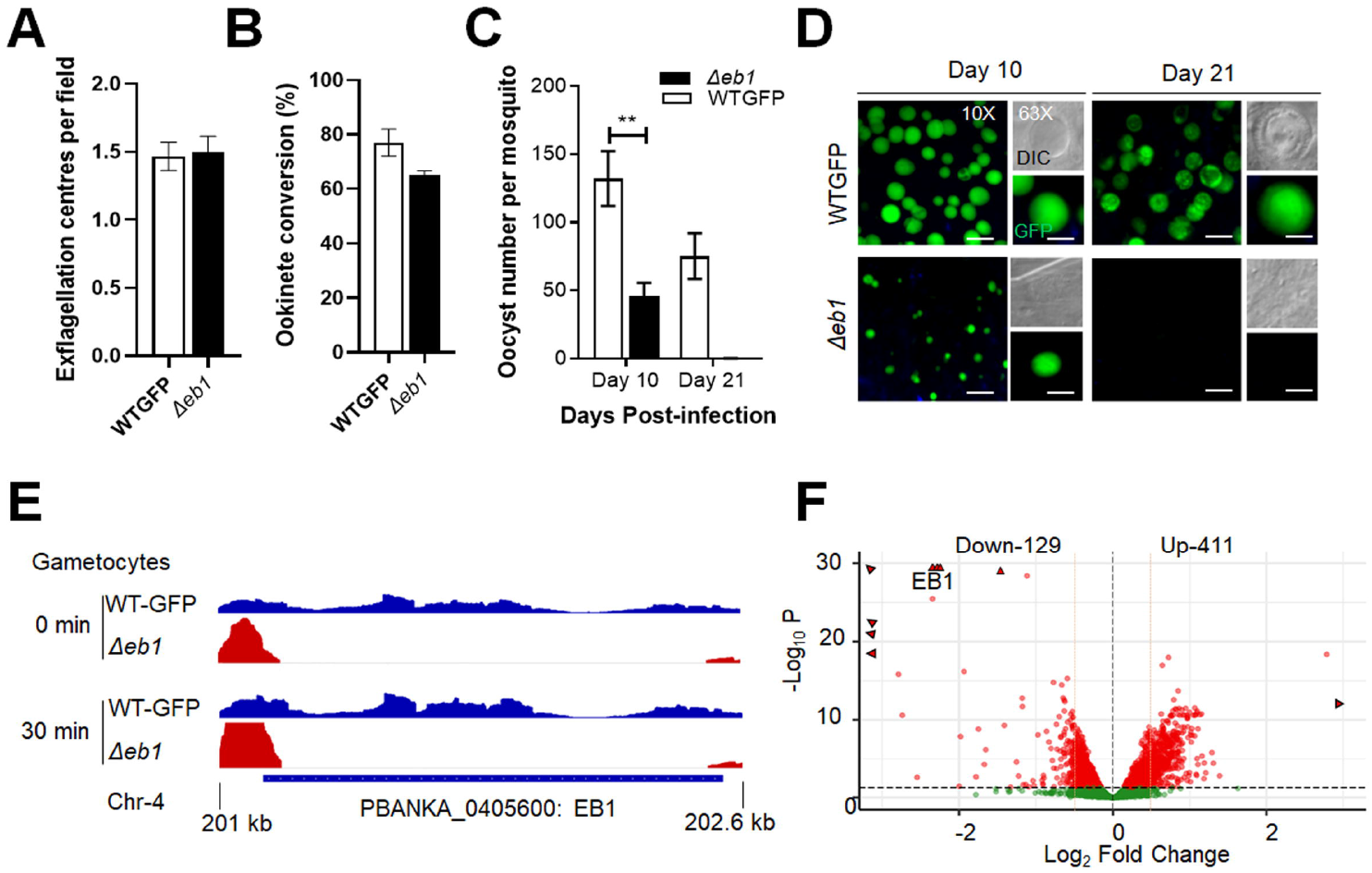
Deletion of *Pbeb1* identifies an essential role in oocyst development and sporogony. **(A)** The number of exflagellation centres per field of Δ*eb1* (black bar) compared with WT-GFP (white bar) parasites at the end of male gametogony. Shown is mean ± SD n = 3 independent experiments. **(B)** Percentage ookinete conversion for Δ*eb1* (black bar) and WT-GFP (white bar) parasites. Ookinetes were identified by reactivity with 13.1 antibody and successful differentiation into elongated ‘banana shaped’ ookinetes. Shown is mean ± SD; n = 3 independent experiments. **(C)** Total number of GFP positive oocysts per infected mosquito in Δ*eb1* (black bar) and WT-GFP (white bar) parasites at 10- and 21-days post-infection (dpi). Shown is mean ± SD; n = 3 independent experiments (with >15 mosquitoes for each) **p<0.01. **(D)** Mid guts at 10x- and 63x-magnification showing fluorescent oocysts of Δ*eb1* and WT-GFP lines at 10 and 21 dpi. Scale bar = 50 μm (10x) or 20 μm (63x). **(E)** RNA-seq analysis showing depletion of EB1 transcript in Δ*eb1* gametocytes at 0- and 30 min post activation. **(F)** Scatter plot showing up- and downregulated genes in Δ*eb1* compared to WT-GFP gametocytes.

Finally, to determine the global pattern of transcription in Δ*eb1* gametocytes we performed RNA-seq analysis 30 minutes after activation. This analysis revealed that, in addition to the complete absence of EB1 transcripts (**Fig 7E**), 129 and 411 genes were significantly downregulated and upregulated respectively (**Fig 7F and Table S4**). GO enrichment analysis of upregulated genes identified proteins involved in phosphorylation, transcription, and microtubule movement **(Fig S10C)**.

### EB1-GFP protein pulldown identifies EB1-MISFIT-MyoK as a putative anchoring complex for ARK2 on spindle MTs

To further analyse the putative interaction of ARK2-GFP with EB1, we performed a reciprocal immunoprecipitation of EB1-GFP, from lysates of paraformaldehyde cross-linked gametocytes one-minute post activation (**Fig 8A, Table S3**). Comparative proteomic analysis of the identified proteins (EB1-GFP versus GFP alone) revealed a pattern of putative interactions for EB1 very similar to that of ARK2, in particular many peptides derived from myosin K and MISFIT. These results suggest that a complex or multiple interactions between EB1, MISFIT, myosin K and ARK2 are present in male gametocytes one minute post activation (**Fig 8A**). Other peptides were derived from components of the kinetochore including INCENP2, ARK1, Stu2 and AKiT1^KNL1^, suggesting that EB1 associates with the spindle-kinetochore interface in a similar way to ARK2. Lastly, we found a strong enrichment of SMC proteins in comparison with GFP alone and ARK2-GFP precipitates, including condensin (SMC2/4) and cohesin (SMC1/3) components, as well as proteins involved in DNA replication (MCM, ORC and RFC).

**Fig 8.**
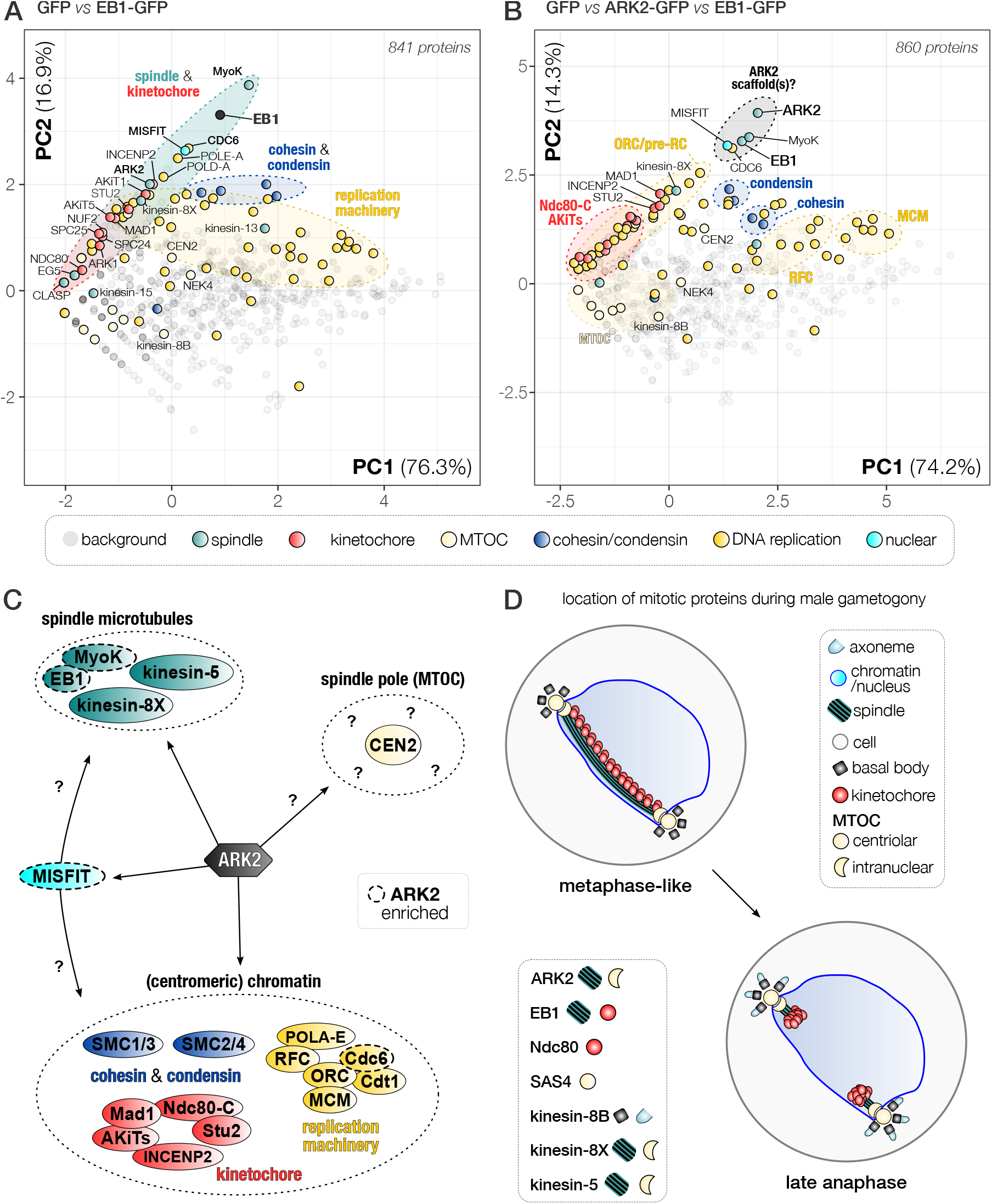
EB1 and ARK2 form part of a protein axis at the spindle during male gametogony. **(A)** Projection of the first two components of a principal component analysis (PCA) of unique peptides identified by mass spectrometry from EB1-GFP or GFP-alone control immunoprecipitates. **(B)** Projection of the first two components of a PCA of unique peptides identified by mass spectrometry from ARK2-GFP, EB1-GFP and GFP-alone immunoprecipitates. Clusters of proteins identified indicate physical and/or functional association, e.g. the MCM (helicase), RFC (replication factor C), condensing (SMC2/4), cohesion (SMC1/3), ORC (Origin of Recognition Complex), parts of the kinetochore and the EB1-ARK2-related putative protein complex (black circle). (**C)** Schematic that reconciles the ARK2 location relative to that of other relevant proteins present at the spindle (pole) during male gametogony. Proteins are grouped by cellular structure/complexes (see legend in panel A). A dashed line indicates proteins that appear enriched in ARK2 and EB1 pulldowns. Bottom: table with numbers of unique peptides in LC-MS/MS analysis of ARK2-GFP and EB1-GFP immunoprecipitates. **(D)** Location of different mitotic proteins during male gametogony. Shown are two phases of mitosis: a metaphase-like state (left top) with kinetochores populating the full length of the spindle, and late anaphase (right bottom).

To further assess whether ARK2, EB1, MISFIT and MyoK associate during male gametogony, we compared ARK2-GFP and EB1-GFP immunoprecipitates.

We reasoned those proteins that would be part of the same protein complex and/or cellular structure should show subtle yet clear co-variation amongst the different PbARK2/EB1-GFP and GFP-only immuno-pulldowns. We therefore used PCA of the combined datasets of unique peptide spectral counts per protein for GFP, ARK2-GFP and EB1-GFP pulldowns (**Fig 8B**). Using the ln(x)+1 transformed peptide values (with non-detected peptide values set to 0), our PCA captures 88.5 % of the variation amongst the data in the first two principal components (see for other principal components **Table S3**). We also found clear clustering for *Pb*ARK2, MISFIT, EB1 and MyoK, consistent with their close association (**Fig 8B**) and a similar pattern for ARK2 and EB1 with the pre-replication complex component Cdc6 (PBANKA_1102900). Furthermore, we observed co-variation for Mad1 (AKiT7), kinesin-8X, the RFC-like protein (PBANKA_0202500) and two polymerase subunits (**Fig 8B**). Overall, we found co-variation in the PCA projection of the ARK2/EB1-GFP pulldown data of proteins that are likely part of the same cellular structures or complexes, such as the kinetochore, spindle and various complexes involved in DNA replication (MCM/RFC/ORC), and the cohesin (SMC1/3) and condensin (SMC2/4) complexes (**Fig 8B-C**), providing further confidence for the value of using PCA for accurate detection of protein complexes. ARK2 and EB1 appear to be transiently part of some of these complexes during male gametogony.

In conclusion, results obtained with our biochemical experiments, in combination with functional analyses, suggest that ARK2 and EB1 are part of a regulatory axis that likely includes MyoK and MISFIT located at, or near the spindle MT-kinetochore interface and are likely involved in the rapid cycles of spindle assembly and chromosome segregation during male gametogony.

## Discussion

Aurora is a serine-threonine kinase family that is highly conserved in eukaryotes. Previous phylogenetic analyses had shown that the family evolved from a single ancestral kinase by widespread recurrent duplications throughout eukaryotic evolution (Willems et al., 2018). AKs play crucial roles in mitotic/meiotic entry, bipolar spindle assembly, chromosome segregation and cytokinesis; and work in conjunction with scaffold proteins like chromosome passenger protein (Hadders and Lens, 2022; Tang et al., 2017; Willems et al., 2018). The three divergent *Plasmodium* AKs are essential for asexual parasite proliferation in the mammalian host but their role in sexual stages and the presence or absence of scaffold proteins were unknown (Solyakov et al., 2011; Tewari et al., 2010). Here we focus on the location and function of *P. berghei* ARK2, an Aurora related protein and its unique scaffold/activator complex, in association with the end binding MT protein (EB1) during the unconventional mode of cell proliferation, differentiation and division during endomitosis and meiosis of sexual transmission stages within the mosquito host.

Our in-depth bioinformatics analysis confirmed the presence of three divergent ARKs and the absence of many scaffold proteins, corroborating earlier studies showing that *Plasmodium* lacks scaffold components like survivin and borealin (van Hooff et al., 2017). However, similar to what was reported in *Toxoplasma* (Berry et al., 2018) two members of INCENP are present. In human cells Aurora A associates with spindle microtubules and centrosomes while Aurora B is located at centromeres, the spindle and the midbody (Carmena and Earnshaw, 2003; Carmena et al., 2015; Hochegger et al., 2013). In other eukaryotes, similar patterns of sub-functionalisation are found (**Fig 1A-C**), with one paralog termed the equatorial AK (Aurora B in humans) and the other the polar AK (Aurora A in humans). Although it is difficult to assign the conserved AK homologue by similar subcellular location of *Plasmodium* ARK2, it appears that ARK2 is more similar to Aurora A due to its association with spindles and with the acentriolar inner MTOC. In such a scenario, ARK2 is the polar AK. Of the other ARKs in *Plasmodium*, we predict that ARK1 is most likely the equatorial ARK due to its conventional AK length (**Fig 1C**) and the association of its one-to-one *T. gondii* ARK1 ortholog with INCENP1-2 (Berry et al., 2018). The presence of a third AK in *Plasmodium* suggests either an additional sub-functionalisation of the canonical equatorial/polar AKs or a new function may have been adopted by ARK3. For both ARK1 and ARK3, similar experiments as performed here need to be conducted to reveal their interactions and functions in the process of chromosome segregation and cell division in *Plasmodium*.

Using live cell imaging of male gametocytes, we show a very discrete and dynamic pattern of ARK2 location during mitosis. The protein transitions from a diffuse nuclear distribution before gametocyte activation to a location at the spindle poles and then moves with the spindle during the three mitotic cycles. A similar pattern is also observed during the first meiotic stages in the developing ookinete. There is no clear anaphase observed in these cells. Live cell imaging of dual fluorescent lines expressing ARK2-GFP and either the kinetochore marker NDC80-mCherry or basal body marker Kinesin8B-mCherry demonstrates that ARK2 occupies a unique location, associated with both spindle MTs and the kinetochore during spindle formation, and is also located at the spindle pole at the inner acentriolar MTOC, but not at the cytoplasmic centriolar MTOC that includes the basal body marker SAS4 or Kinesin8B (Rashpa and Brochet, 2022; Zeeshan et al., 2022). STED microscopy with alpha-tubulin and super-resolution images of dual fluorescence-tagged lines further corroborate this unusual location of ARK2. The number of acentriolar MTOCs, defined by the location of ARK2-GFP during male gamete formation and zygote differentiation correlates with the ploidy of the cell, for example in the 2N, 4N or 8N male cell there were two, four or eight foci, and in the 4N ookinete there were four foci. Interestingly the ARK2-GFP signal disappeared by 12 hours of zygote development, when it is likely that the second meiotic division had taken place (without karyokinesis), whereas NDC80 was present until the end of the ookinete stage but in both cases four fluorescent foci are seen in the 4N ookinete.

EB1 is a plus-end MT tracking protein that accumulates at the growing ends of MTs and has a key role in the regulation of MT dynamics (Komarova et al., 2009). During male gametogony, EB1 had a location similar to that of ARK2 on the spindle and acentriolar MTOC. The parasite line expressing dual fluorescent-tagged EB1 and ARK2 showed that they are closely associated with each other at the different stages of development during both male gamete formation and zygote differentiation. STED imaging of EB1 showed that EB1 is associated with the spindle as was also observed for ARK2, suggesting that both are binding to spindle MTs. Intriguingly, EB1 was not detected in the proliferative asexual stages within red blood cells. This is in contrast to the presence of EB1 during the non-mitotic gametocytogenesis in *P. falciparum* (Li et al., 2022) and in asexual cell proliferation in *Toxoplasma* where EB1 was observed associated with spindle MTs (Chen et al., 2015). These findings suggest that both ARK2 and EB1 may be a part of the spindle machinery and the acentriolar MTOC during male gamete formation and ookinete development.

Our previous *Plasmodium* kinome screen showed that ARK2 has an essential role during blood stage development (Tewari et al., 2010). Here we used conditional knockdown approaches to study the functional role of ARK2 in sexual stages and our results show that our *ark2* and *eb1* knockdown mutants have a similar phenotype to those of *Plasmodium*-specific cyclin, PbCYC3, and kinesin-8X, in which oocyst size and sporozoite formation were affected (Roques et al., 2015; Zeeshan et al., 2019b), and similar to what is observed in other deletion mutants including MISFIT, PK7 and PPM5 genes (Bushell et al., 2009; Guttery et al., 2014; Tewari et al., 2010). Genetic backcross experiments with *dozi* and *nek4* mutants that affect female and male gametogony, respectively, demonstrated that the *P*_*clag-*_*ark2* defect in oocyst development is inherited as a defect in the male gametocyte lineage, similar to what is observed for Δ*misfit* and Δ*ppm5*, for which there is an absolute requirement for a functional gene from the male line (Bushell et al., 2009; Guttery et al., 2014). These data suggest that both ARK2 and EB1 are part of the spindle assembly, and although male gametes and ookinetes are produced, downregulated ARK2 expression has a delayed effect that is seen during oocyst development and results in a complete block in parasite transmission.

Global transcript analysis showed significant differences in gene expression between the knockdown *P*_*clag-*_*ark2* parasites and WT lines. Genes coding for proteins involved in MT-based movement and regulation of gene expression were mostly affected, including a large number of protein kinases; several motor proteins (e.g. kinesin and dynein); and proteins involved in invasion or oocyst development. This finding is consistent with global phospho-proteomic studies of male gametogony, in which ARK2 was shown to be associated with rapid phosphorylation of MT proteins in either very early or late stages of male gamete formation (Invergo et al., 2017). It is possible that ARK2 phosphorylates various substrates including kinesin-8X, EB1, and MISFIT. In all these cases the genetic defect is transferred through the male lineage and manifest during endomitosis in the oocyst; thereby blocking parasite development and transmission.

Our results suggest that *Pb*ARK2 is largely localised on the spindle apparatus associated with kinetochores, suggesting that it is not part of a CPC-like complex. We confirmed the earlier phylogenetic studies that showed that CPC components like Survivin and Borealin are absent and ARK2-GFP immunoprecipitations identified unique candidate ARK2-interacting proteins. Kinetochore components and proteins with a role at the spindle apparatus were identified. These proteins included the MT plus-end binding protein EB1, the myosin MyoK, a nuclear formin-like protein called MISFIT, members of the NDC80 outer kinetochore complex, and other Apicomplexan Kinetochore proteins (AKiTs). The presence of such interactors strongly suggests that ARK2 binds in proximity to the kinetochore-spindle MT interface. A reciprocal pulldown with EB1-GFP identified a similar set of interacting proteins as ARK2-GFP. Components of the kinetochore like MAD1, NDC80, STU2 and AkiT were detected although no high abundance peptides were present in both ARK2 and EB1 pulldowns or highlighted by PCA. In addition, none of the TPX2 complex components were detected, suggesting that ARK2 may not be exactly functionally similar to Aurora A (polar) of model eukaryotes (Willems et al., 2018). This presence of a unique plasmodium ARK2 scaffold protein and its localisation suggest that it may have a cross-functional role in relation to conventional Aurora A and Aurora B.

*Plasmodium* has only one EB1 homologue, compared to the three EB1 proteins that exist in other eukaryotes (Komarova et al., 2009). EB1 function is heavily regulated by protein phosphorylation: a cluster of six serine residues present in the linker region of the yeast EB1 homologue (Bim1) is phosphorylated by the AK homologue lpl1, regulating disassembly of the spindle midzone during anaphase. Human EB1 is co-immunoprecipitated with Aurora B (Sun et al., 2008); the EB1 concentrates Aurora B at inner centromeres in a MT-dependent manner, resulting in phosphorylation of both kinetochore and chromatin substrates (Banerjee et al., 2014).

The interaction of ARK2 with EB1, MyoK and Misfit was revealed through co-variation of these proteins in ARK2 and EB1-GFP immunoprecipitates as measured using PCA. These data suggest that ARK2 in *Plasmodium* may form a unique complex with these proteins that has not been described in other organisms. MyoK has been shown in many studies to be involved in mitosis in many organisms however MISFIT is a *Plasmodium* specific formin (Bushell et al., 2009). How ARK2 interacts with EB1, MyoK and Misfit is unclear. Possibly its extended length, with the presence of a long-coiled coil in *Plasmodium* spp. (Fig 1C) facilitates interactions with other long coiled regions, such as found in MyoK for instance. It not known whether MyoK and Misfit are part of the spindle MT as seen for EB1, and this will need to be investigated in future studies. The presence of this unique association of these scaffold proteins suggest that it may be related to an unconventional mode of lateral spindle apparatus and chromosome segregation that is observed in these parasite sexual stages.

Overall, this study suggests that *Plasmodium* ARK2 is an Aurora paralogue that is located at the spindle and spindle poles formed by the acentriolar MTOC. It forms a unique association with EB1 and some kinetochore molecules but not in a way similar to Aurora B, which is CPC based (INCENP/Borealin/Survivin), nor Aurora A (TPX/Cep192/ BORA). Hence ARK2 uniquely interacts with a putative Aurora scaffold consisting of EB1/MISFIT/MyoK that is highly divergent compared to other eukaryotes, and that drives endomitosis and meiosis during parasite transmission. This suggests the flexibility of molecular networks to rewire and drive unconventional modes of spindle organisation and chromosome segregation during cell division in the malaria parasite *Plasmodium*.

## Materials and Methods

### Ethics statement

The animal work passed an ethical review process and was approved by the United Kingdom Home Office. Work was carried out under UK Home Office Project Licenses (30/3248 and PDD2D5182) in accordance with the UK ‘Animals (Scientific Procedures) Act 1986’. Six- to eight-week-old female CD1 outbred mice from Charles River laboratories were used for all experiments.

### Generation of transgenic parasites and genotype analyses

To generate the GFP-tag lines, a region of each gene (*ark2 and eb1*) downstream of the ATG start codon was amplified, ligated to p277 vector, and transfected as described previously (Guttery et al., 2012). The p277 vector contains the human *dhfr* cassette, conveying resistance to pyrimethamine. A schematic representation of the endogenous gene locus, the constructs and the recombined gene locus can be found in **Fig S1A and S5A**. For the parasites expressing the C-terminal GFP-tagged protein, diagnostic PCR was used with primer 1 (Int primer) and primer 3 (ol492) to confirm integration of the GFP targeting construct **(Fig S1B and S5B)**. A list of primers used to amplify these genes can be found in **Table S5**.

For the generation of transgenic *ark2*-AID/HA line, library clone PbG01-2471h08 from the PlasmoGEM repository (http://plasmogem.sanger.ac.uk/) was used. Sequential recombineering and gateway (GW) steps were performed as previously described (Pfander et al., 2013; Pfander et al., 2011). Insertion of the GW cassette following gateway reaction was confirmed using primer pairs GW1 (CATACTAGCCATTTTATGTG) x *ark2* QCR1 (GCTTTGCAGCCGAAGCTCCG) and GW2 (CTTTGGTGACAGATACTAC) x *ark2* QCR2 (AGGGGGAAAATGTTACACATGCGT). The modified library inserts were then released from the plasmid backbone using *Not*I. The *ark2*-AID/HA targeting vector was transfected into the 615-parasite line and conditional degradation of ARK-AID/HA in the non-clonal line was performed as described previously (Balestra et al., 2020). A schematic representation of the endogenous *ark2* locus, the constructs and the recombined *ark2* locus can be found in **Fig S3A**. A diagnostic PCR was performed for *ark2* gene knockdown parasites as outlined in **Fig S3A**. Primer pairs *ark2* QCR1/GW1, and *ark2* QCR2/GW2 were used to determine successful integration of the targeting construct at the 3’ end of the gene **(Fig S3B)**.

The conditional knockdown construct *P*_*clag-*_*ark2* was derived from *P*_*clag*_ (pSS367) by placing *ark2* under the control of the *clag* gene (PBANKA_083630) promoter, as described previously (Sebastian et al., 2012). A schematic representation of the endogenous *ark2* locus, the constructs and the recombined *ark2* locus can be found in **Fig S3E**. A diagnostic PCR was performed for *ark2* gene knockdown parasites as outlined in **Fig S3E**. Primer 1 (5’-intPTD24) and Primer 2 (5’-intPTD) were used to determine successful integration of the targeting construct at the 5’ end of the gene. Primer 3 (3’-intPTclag) and Primer 4 (3’-intPTD24) were used to determine successful integration for the 3’ end of the gene locus **(Fig S3F)**. All the primer sequences can be found in **Table S5**.

To study the function of EB1, the gene-deletion targeting vector for *eb1 was* constructed using the pBS-DHFR plasmid, which contains polylinker sites flanking a *T. gondii dhfr/ts* expression cassette conferring resistance to pyrimethamine, as described previously (Tewari et al., 2010). The 5′ upstream sequence of *eb1* was amplified from genomic DNA and inserted into *Apa*I and *Hin*dIII restriction sites upstream of the *dhfr/ts* cassette of pBS-DHFR. A DNA fragment amplified from the 3′ flanking region of *eb1* was then inserted downstream of the *dhfr/ts* cassette using *Eco*RI and *Xba*I restriction sites. The linear targeting sequence was released using *Apa*I/*Xba*I. A schematic representation of the endogenous *eb1* locus, the construct and the recombined *eb1* locus can be found in **Fig S7A**. The primers used to generate the mutant parasite lines can be found in **Table S5**. A diagnostic PCR was used with primer 1 (IntN138_5) and primer 2 (ol248) to confirm integration of the targeting construct, and primer 3 (KO1) and primer 4 (KO2) were used to confirm deletion of the *eb1* gene **(Fig S7B, Table S5)**. *P. berghei* ANKA line 2.34 (for GFP-tagging) or ANKA line 507cl1 expressing GFP (for the gene deletion and knockdown construct) parasites were transfected by electroporation (Janse et al., 2006).

### Live cell imaging

To examine ARK2-GFP and EB1-GFP expression during erythrocytic stages, parasites growing in schizont culture medium were used for imaging at different stages of schizogony. Purified gametocytes were examined for GFP expression and cellular location at different time points (0, 1-15 min) after activation in ookinete medium (Zeeshan et al., 2019b). Zygote and ookinete stages were analysed throughout 24 h of culture using cy3-conjugated 13.1 antibody (red), which recognises P28 protein on the surface of zygotes and ookinetes. Oocysts and sporozoites were imaged using infected mosquito guts. Images were captured using a 63x oil immersion objective on a Zeiss Axio Imager M2 microscope fitted with an AxioCam ICc1 digital camera.

### Generation of dual tagged parasite lines

The green (GFP)- or red (mCherry)-tagged ARK2 and EB1 parasite lines were mixed with mCherry- or GFP-tagged lines of kinetochore marker NDC80 (Zeeshan et al., 2020b), axoneme marker kinesin-8B (Zeeshan et al., 2019a) and basal body marker SAS4 (Zeeshan et al., 2022) in equal numbers and injected into mice. Mosquitoes were fed on these mice 4 to 5 days after infection when gametocytemia was high, and were checked for oocyst development and sporozoite formation at day 14 and day 21 after feeding. Infected mosquitoes were then allowed to feed on naïve mice and after 4 to 5 days the mice were examined for blood stage parasitaemia by microscopy with Giemsa-stained blood smears. Some parasites expressed both ARK2-mCherry and NDC80-GFP; and ARK2-GFP and kinesin-8B-cherry in the resultant gametocytes, and these were purified, and fluorescence microscopy images were collected as described above.

### Parasite phenotype analyses

Blood samples containing approximately 50,000 parasites of the *ark2* knockdown/*eb1* knockout lines were injected intraperitoneally (i.p) into mice to initiate infection. Asexual stages and gametocyte production were monitored by microscopy on Giemsa-stained thin smears. Four to five days post infection, exflagellation and ookinete conversion were examined as described previously (Guttery et al., 2012) with a Zeiss AxioImager M2 microscope (Carl Zeiss, Inc) fitted with an AxioCam ICc1 digital camera. To analyse mosquito infection and transmission, 30 to 50 *Anopheles stephensi* SD 500 mosquitoes were allowed to feed for 20 min on anaesthetized, infected mice with at least 15% asexual parasitaemia and carrying comparable numbers of gametocytes as determined on Giemsa-stained blood films. To assess mid-gut infection, approximately 15 guts were dissected from mosquitoes on days 7 and 14 post feeding and oocysts were counted using a 63x oil immersion objective. On day 21 post-feeding, another 20 mosquitoes were dissected, and their guts and salivary glands crushed separately in a loosely fitting homogenizer to release sporozoites, which were then quantified using a haemocytometer or used for imaging. Mosquito bite-back experiments were performed 21 days post-feeding using naive mice, and blood smears were examined after 3-4 days.

### Purification of gametocytes

The purification of gametocytes was achieved by injecting parasites into phenylhydrazine treated mice (Beetsma et al., 1998) and gametocyte enrichment by sulfadiazine treatment after 2 days of infection. The blood was collected on day 4 after infection and gametocyte-infected cells were purified on a 48% v/v NycoDenz (in PBS) gradient (NycoDenz stock solution: 27.6% w/v NycoDenz in 5 mM Tris-HCl, pH 7.20, 3 mM KCl, 0.3 mM EDTA). The gametocytes were harvested from the interface and activated.

### Immunoprecipitation and mass spectrometry

Male gametocytes of ARK2-GFP and EB1-GFP parasites were used at 1 min post activation to prepare cell lysates. WT-GFP gametocytes were used as controls. Purified parasite pellets were crosslinked using formaldehyde (10□min incubation with 1% formaldehyde, followed by 5□min incubation in 0.125□M glycine solution and three washes with phosphate-buffered saline (PBS; pH 7.5). Immunoprecipitation was performed using the protein lysates and a GFP-Trap_A Kit (Chromotek) following the manufacturer’s instructions. Briefly, the lysates were incubated for 2h with GFP-Trap_A beads at 4º C with continuous rotation. Unbound proteins were washed away, and proteins bound to the GFP-Trap_A beads were digested using trypsin. The tryptic peptides were analysed by liquid chromatography–tandem mass spectrometry. Mascot (http://www.matrixscience.com/) and MaxQuant (https://www.maxquant.org/) search engines were used for mass spectrometry data analysis. Peptide and proteins having a minimum threshold of 95% were used for further proteomic analysis. The PlasmoDB database was used for protein annotation, and a separate manual curation was performed to classify proteins into 6 categories relevant for functional annotation of ARK2/EB1 immunoprecipitates: background, cohesin/condensin, DNA repair/replication, kinetochore, MTOC, spindle, proteasome and ribosome/translation (Metsalu and Vilo, 2015). The first six principal components for the analysis comparing ARK2/EB1/GFP-only samples can be found in **Table S3**.

### Ookinete motility assays

The motility of *P*_*clag-*_*ark2* ookinetes was assessed using Matrigel as described previously (Volkmann et al., 2012; Zeeshan et al., 2020a). Ookinete cultures grown for 24 h were added to an equal volume of Matrigel (Corning), mixed thoroughly, dropped onto a slide, covered with a cover slip, and sealed with nail polish. The Matrigel was then allowed to set at 20°C for 30 min. After identifying a field containing an ookinete, time-lapse videos (one frame every 5 s for 100 cycles) were collected using the differential interference contrast settings with a 63× objective lens on a Zeiss AxioImager M2 microscope fitted with an AxioCam ICc1 digital camera and analysed with the AxioVision 4.8.2 software.

### Fixed immunofluorescence assay and deconvolution microscopy

The ARK2-GFP gametocytes were purified, activated in ookinete medium, fixed at different time points with 4% paraformaldehyde (PFA, Sigma) diluted in MT-stabilising buffer (MTSB) for 10-15 min, and added to poly-L-lysine coated slides. Immunocytochemistry was performed using primary GFP-specific rabbit monoclonal antibody (mAb) (Invitrogen-A1122; used at 1:250) and primary mouse anti-α tubulin mAb (Sigma-T9026; used at 1:1000). Secondary antibodies were Alexa 488 conjugated anti-mouse IgG (Invitrogen-A11004) and Alexa 568 conjugated anti-rabbit IgG (Invitrogen-A11034) (used at 1 in 1000). The slides were then mounted in Vectashield 19 with DAPI (Vector Labs) for fluorescence microscopy. Parasites were visualised on a Zeiss AxioImager M2 microscope fitted with an AxioCam ICc1 digital camera. Post-acquisition analysis was carried out using Icy software – version 1.9.10.0. Images presented are 2D projections of deconvoluted Z-stacks of 0.3□μm optical sections.

### STED microscopy

Immunofluorescence staining for STED microscopy was performed as a combination of protocols described previously (Ponjavic et al., 2021; Simon et al., 2021). Briefly, the gametocytes were fixed with 4% pre-warmed PFA/PBS. PFA was washed away thrice with PBS. Fixed cells were stored in PBS at 4°C in the dark for later immunofluorescence staining. Before beginning the immunofluorescence procedure, glass-coated 35 mm imaging μ-dishes (Ibidi, 81156) were coated with poly-L-lysine (PLL) solution (0.01%, Sigma-Aldrich, P4832) according to the manufacturer’s guidelines (0.05% final solution). After extensive washing with nuclease-free water, dishes were left to dry. The fixed cells in PBS were then seeded into PLL-coated dishes and left to settle for a day. The cells were then washed with PBS, permeabilized with 0.5% Triton X-100/PBS for 30 min at room temperature and rinsed three times with PBS. To quench free aldehyde groups, cells were incubated with freshly prepared 0.1 mg/ml NaBH4/PBS solution for 10 min. Cells were rinsed thrice with PBS and blocked with 3% BSA/PBS for 30 min. In the meantime, primary antibodies were diluted in 3% BSA/PBS and the solution was centrifuged at 21,100*g* for 10 min at 4°C to remove potential aggregates. Cells were incubated with primary antibody to stain tubulin (mouse anti-α-tubulin B-5-1-2 mAb, Sigma-Aldrich, T5168, dilution 1:250) for 4 h at room temperature. Next, the cells were washed three times with 0.5% Tween-20/PBS. Incubation with secondary antibodies (donkey anti-mouse IgG Alexa Fluor 594, Abcam, ab150112; RRID: AB_2813898, dilution 1:500 or STAR ORANGE, goat anti-mouse IgG, Abberior GmbH, STORANGE-1001-500UG, dilution 1:500) in 3% BSA/PBS was performed for 1 h after removal of aggregates as described for primary antibodies. After washing twice with 0.5% Tween-20/PBS and once with PBS, cells were incubated with SiR-DNA solution (Spirochrome, SC007, 1:100) for 1h, then washed once washed with PBS and stored in PBS at 4°C in the dark until imaging.

Rescue-STED microscopy was performed on a single-point scanning Expert Line easy3D STED super-resolution microscope (Abberior Instruments GmbH), equipped with a pulsed 775 nm STED depletion laser and two avalanche photodiodes for detection. Super-resolution images were acquired with a 100×1.4 NA objective, a pixel size of 10-20 nm and a pixel dwell time of 8 µs. The STED laser power was set to 15–30%, whereas the other lasers (488, 594 and 640 nm) were adjusted to the antibody combinations used. To acquire z-stacks, a total z-stack of 3-5 μm was acquired using a z-step size of 200-300 nm. The channel where the 640 nm laser was used for SiR-DNA excitation was taken separately in time but in the same imaging region as used for the 488 and 594 channels, and with custom-made defined emission boundaries of 594 and 640 to limit signal crosstalk between the channels. 640 and 594 nm channels were taken with STED depletion laser using the parameters described above, whereas 488 channel was taken with the same parameters but without STED depletion laser. STED images were assembled in Fiji (ImageJ-win64) as maximum intensity projections of acquired z-stacks that contained noticeable EB1 and ARK2 signals.

### Ultrastructure expansion microscopy (U-ExM)

Purified gametocytes were activated for 1-2 minutes and then activation was stopped by adding 4% formaldehyde. Sample preparation of *P. berghei* parasites for U-ExM was performed as previously described (Bertiaux et al., 2021; Gambarotto et al., 2021), except that 4% formaldehyde (FA) was used as fixative (Rashpa and Brochet, 2022). Fixed samples were then attached on a 12 mm round Poly-D-Lysine (A3890401, Gibco) coated coverslips for 10 minutes. Immuno-labelling was performed using primary antibodies against α-tubulin and β-tubulin (1:200 dilution, AA344 and AA345 from the Geneva antibody facility), anti γ-tubulin antibody (1:500 dilution, Sigma T5192) and anti HA antibody (3F10) (1:250 dilution, Roche). Secondary antibodies anti-guinea pig Alexa 647, anti-rabbit Alexa 405 and anti-rat Alexa 488 were used at dilutions 1:400 (Invitrogen). Atto 594 NHS-ester was used for bulk proteome labelling (Merck 08741). Images were acquired on a Leica TCS SP8 microscope, image analysis was performed using Fiji-Image J and Leica Application Suite X (LAS X) software.

### Structured illumination microscopy

A small volume (3 µl) of gametocytes was mixed with Hoechst dye and pipetted onto

2 % agarose pads (5×5 mm squares) at room temperature. After 3 min these agarose pads were placed onto glass bottom dishes with the cells facing towards glass surface (MatTek, P35G-1.5-20-C). Cells were scanned with an inverted microscope using Zeiss C-Apochromat 63×/1.2 W Korr M27 water immersion objective on a Zeiss Elyra PS.1 microscope, using the structured illumination microscopy (SIM) technique. The correction collar of the objective was set to 0.17 for optimum contrast. The following settings were used in SIM mode: lasers, 405 nm: 20%, 488 nm: 50%; exposure times 100 ms (Hoechst) and 25 ms (GFP); three grid rotations, five phases. The band pass filters BP 420-480 + LP 750 and BP 495-550 + LP 750 were used for the blue and green channels, respectively. Multiple focal planes (Z stacks) were recorded with 0.2 µm step size; later post-processing, a Z correction was done digitally on the 3D rendered images to reduce the effect of spherical aberration (reducing the elongated view in Z; a process previously tested with fluorescent beads). Images were processed and all focal planes were digitally merged into a single plane (Maximum intensity projection). The images recorded in multiple focal planes (Z-stack) were 3D rendered into virtual models and exported as images and movies (see supplementary material). Processing and export of images and videos were done by Zeiss Zen 2012 Black edition, Service Pack 5 and Zeiss Zen 2.1 Blue edition.

### RNA isolation and quantitative Real Time PCR (qRT-PCR) analyses

RNA was isolated from purified gametocytes using an RNA purification kit (Stratagene). cDNA was synthesized using an RNA-to-cDNA kit (Applied Biosystems). Gene expression was quantified from 80 ng of total RNA using SYBR green fast master mix kit (Applied Biosystems). All the primers were designed using primer3 (Primer-blast, NCBI). Analysis was conducted using an Applied Biosystems 7500 fast machine with the following cycling conditions: 95°C for 20 s followed by 40 cycles of 95°C for 3 s; 60°C for 30 s. Three technical replicates and three biological replicates were performed for each assayed gene. The *hsp70* (PBANKA_081890) and *arginyl-t RNA synthetase* (PBANKA_143420) genes were used as endogenous control reference genes. The primers used for qPCR can be found in **Table S5**.

### RNA-seq analysis

Libraries were prepared from lyophilized total RNA, first by isolating mRNA using NEBNext Poly(A) mRNA Magnetic Isolation Module (NEB), then using NEBNext Ultra Directional RNA Library Prep Kit (NEB) according to the manufacturer’s instructions. Libraries were amplified for a total of 12 PCR cycles (12 cycles of [15 s at 98°C, 30 s at 55°C, 30 s at 62°C]) using the KAPA HiFi HotStart Ready Mix (KAPA Biosystems). Libraries were sequenced using a NovaSeq 6000 DNA sequencer (Illumina), producing paired-end 100-bp reads.

FastQC (https://www.bioinformatics.babraham.ac.uk/projects/fastqc/), was used to analyse raw read quality. The first 11 bp of each read and any adapter sequences were removed using Trimmomatic (http://www.usadellab.org/cms/?page=trimmomatic). Bases were trimmed from reads using Sickle with a Phred quality threshold of 25 (https://github.com/najoshi/sickle). The resulting reads were mapped against the *P. berghei* ANKA genome (v36) using HISAT2 (version 2-2.1.0), using default parameters. Uniquely mapped, properly paired reads with mapping quality 40 or higher were retained using SAMtools (http://samtools.sourceforge.net/). Genome browser tracks were generated and viewed using the Integrative Genomic Viewer (IGV) (Broad Institute). Raw read counts were determined for each gene in the *P. berghei* genome using BedTools (https://bedtools.readthedocs.io/en/latest/#) to intersect the aligned reads with the genome annotation. Differential expression analysis was done by use of R package DESeq2 to call up- and down-regulated genes with an adjusted P-value cutoff of 0.05. Gene ontology enrichment was done using R package topGO (https://bioconductor.org/packages/release/bioc/html/topGO.html) with the weight01 algorithm.

### ChIP-seq analysis

Gametocytes of EB1-GFP and NDC80-GFP (as a positive control) parasites were harvested, and the pellets were resuspended in 500 µl of Hi-C lysis buffer (25 mM Tris-HCl, pH 8.0, 10 mM NaCl, 2 mM AESBF, 1% NP-40, protease inhibitors). After incubation for 10 min at room temperature (RT), the resuspended pellets were homogenized by passing through a 26.5 gauge needle/syringe 15 times and cross-linked by adding formaldehyde (1.25% final concentration) for 25 min at RT with continuous mixing. Crosslinking was stopped by adding glycine to a final concentration of 150 mM and incubating for 15 min at RT with continuous mixing. The sample was centrifuged for 5 min at 2,500 x g (∼5,000 rpm) at 4°C, the pellet washed once with 500 µl ice-cold wash buffer (50 mM Tris-HCl, pH 8.0, 50 mM NaCl, 1 mM EDTA, 2 mM AESBF, protease inhibitors) and the pellet stored at -80°C for ChIP-seq analysis. The crosslinked parasite pellets were resuspended in 1 mL of nuclear extraction buffer (10 mM HEPES, 10 mM KCl, 0.1 mM EDTA, 0.1 mM EGTA, 1 mM DTT, 0.5 mM AEBSF, 1X protease inhibitor tablet), post 30 min incubation on ice, 0.25% Igepal-CA-630 was added and the sample homogenized by passing through a 26G x ½ needle. The nuclear pellet extracted through 5,000 rpm centrifugation, was resuspended in 130 µl of shearing buffer (0.1% SDS, 1 mM EDTA, 10 mM Tris-HCl pH 7.5, 1X protease inhibitor tablet), and transferred to a 130 µl Covaris sonication microtube. The sample was then sonicated using a Covaris S220 Ultrasonicator for 8 min (Duty cycle: 5%, intensity peak power: 140, cycles per burst: 200, bath temperature: 6°C). The sample was transferred to ChIP dilution buffer (30 mM Tris-HCl pH 8.0, 3 mM EDTA, 0.1% SDS, 30 mM NaCl, 1.8% Triton X-100, 1X protease inhibitor tablet, 1X phosphatase inhibitor tablet) and centrifuged for 10 min at 13,000 rpm at 4°C, retaining the supernatant. For each sample, 13 μl of protein A agarose/salmon sperm DNA beads were washed three times with 500 µl ChIP dilution buffer (without inhibitors) by centrifuging for 1 min at 1,000 rpm at room temperature, then buffer was removed. For pre-clearing, the diluted chromatin samples were added to the beads and incubated for 1 hour at 4°C with rotation, then pelleted by centrifugation for 1 min at 1,000 rpm. Before adding antibody, ∼10% of one EB1-GFP sample was taken as input. Supernatant was removed into a LoBind tube, carefully so as not to remove any beads, and 2 µg of anti-GFP antibody (Abcam ab290, anti-rabbit) were added to the sample and incubated overnight at 4°C with rotation. For one EB1-GFP sample, IgG antibody (ab37415) was added instead as a negative control. Per sample, 25 µl of protein A agarose/salmon sperm DNA beads were washed with ChIP dilution buffer (no inhibitors), blocked with 1 mg/mL BSA for 1 hour at 4°C, then washed three more times with buffer. 25 µl of washed and blocked beads were added to the sample and incubated for 1 hour at 4°C with continuous mixing to collect the antibody/protein complex. Beads were pelleted by centrifugation for 1 min at 1,000 rpm at 4°C. The bead/antibody/protein complex was then washed with rotation using 1 mL of each buffers twice; low salt immune complex wash buffer (1% SDS, 1% Triton X-100, 2 mM EDTA, 20 mM Tris-HCl pH 8.0, 150 mM NaCl), high salt immune complex wash buffer (1% SDS, 1% Triton X-100, 2 mM EDTA, 20 mM Tris-HCl pH 8.0, 500 mM NaCl), high salt immune complex wash buffer (1% SDS, 1% Triton X-100, 2 mM EDTA, 20 mM Tris-HCl pH 8.0, 500 mM NaCl), TE wash buffer (10 mM Tris-HCl pH 8.0, 1 mM EDTA) and eluted from antibody by adding 250 μl of freshly prepared elution buffer (1% SDS, 0.1 M sodium bicarbonate). We added 5 M NaCl to the elution and cross-linking was reversed by heating at 45°C overnight followed by addition of 15 μl of 20 mg/mL RNAase A with 30 min incubation at 37°C. After this, 10 μl 0.5 M EDTA, 20 μl 1 M Tris-HCl pH 7.5, and 2 μl 20 mg/mL proteinase K were added to the elution and incubated for 2 hours at 45°C. DNA was recovered by phenol/chloroform extraction and ethanol precipitation, using a phenol/chloroform/isoamyl alcohol (25:24:1) mixture twice and chloroform once, then adding 1/10 volume of 3 M sodium acetate pH 5.2, 2 volumes of 100% ethanol, and 1/1000 volume of 20 mg/mL glycogen. Precipitation was allowed to occur overnight at -20°C. Samples were centrifuged at 13,000 rpm for 30 min at 4°C, then washed with fresh 80% ethanol, and centrifuged again for 15 min with the same settings. Pellet was air-dried and resuspended in 50 μl nuclease-free water. DNA was purified using Agencourt AMPure XP beads. Libraries were then prepared from this DNA using a KAPA library preparation kit (KK8230) and sequenced on a NovaSeq 6000 machine. FastQC (https://www.bioinformatics.babraham.ac.uk/projects/fastqc/), was used to analyze raw read quality. Any adapter sequences were removed using Trimmomatic (http://www.usadellab.org/cms/?page=trimmomatic). Bases with Phred quality scores below 25 were trimmed using Sickle (https://github.com/najoshi/sickle). The resulting reads were mapped against the *P. berghei* ANKA genome (v36) using Bowtie2 (version 2.3.4.1). Using Samtools, only properly paired reads with mapping quality 40 or higher were retained and reads marked as PCR duplicates were removed by PicardTools MarkDuplicates (Broad Institute). Genome-wide read counts per nucleotide were normalized by dividing millions of mapped reads for each sample (for all samples including input) and subtracting input read counts from the ChIP and IgG counts. From these normalized counts, genome browser tracks were generated and viewed using the Integrative Genomic Viewer (IGV).

### Statistical analysis

All statistical analyses were performed using GraphPad Prism 7 (GraphPad Software). For qRT-PCR, a two-way ANOVA test was used to examine significant differences between wild-type and mutant strains.

## Data Availability

DNA Sequence reads have been deposited in the NCBI Sequence Read Archive with accession number: PRJNA808974

## Acknowledgments

We wish to thank Julie Rodgers for helping to maintain the insectary and other technical works and Cleidiane Zampronio at University Warwick for mass mass spectrometry methods,

## SUPPLEMENTARY DATA

## Supplementary figures

**Fig S1. Generation of PbARK2-GFP parasites and analysis of subcellular location of ARK2-GFP throughout the life cycle**

**(A)** Schematic representation of the endogenous *Pbark2* locus, the GFP-tagging construct and the recombined *ark2* locus following single homologous recombination. Arrows 1 and 2 indicate the position of PCR primers used to confirm successful integration of the construct. **(B)** Diagnostic PCR of *ark2* and WT parasites using primers IntT204 (Arrow 1) and ol492 (Arrow 2). Integration of the ark2 tagging construct gives a band of 594 bp. Tag = ARK2-GFP parasite line. **(C)** Live cell imaging of ARK2-GFP parasites during erythrocytic schizogony showing one or two focal points of ARK2-GFP (green) per nucleus. DNA is stained with Hoechst dye (blue); scale bar = 5 μm. **(D)** Live cell imaging of ARK2-GFP parasites during oocyst development in mosquitoes showing discrete foci of ARK2-GFP. DNA is stained with Hoechst dye (blue); scale bar = 5 μm. **(E)** Live cell imaging showing ARK2-GFP gametocytes at 30 sec and 15 min after activation. ARK2-GFP was not detected in free gametes (15 min gametocytes). DNA is stained with Hoechst dye (blue); scale bar = 5 μm. **(F)** Live-cell imaging showing ARK2-GFP location in zygote and ookinete. A cy3-conjugated antibody, 13.1, which recognises the protein P28 on the surface of zygotes and ookinetes was used to mark these stages (red). DNA is stained with Hoechst dye (blue); scale bar = 5 μm. **(G)** Still images (at every 5 s) showing dynamic location of ARK2-GFP in gametocytes within 3 to 4 min post activation (mpa) during male gametogony. DNA is stained with Hoechst dye (blue); scale bar = 5 μm. **(H)** Still images (at every 5 s) showing dynamic location of ARK2-GFP within 6 to 7 mpa during male gametogony. DNA is stained with Hoechst dye (blue); scale bar = 5 μm.

**Fig S2. Quantification and staining with tubulin antibody of events of ARK2 localization during male gametogony**.

**A**. The events of ARK2-GFP localization during different time points after gametocytes activation. **(B)** Immunofluorescence assay (IFA) showing location of ARK2 (green) and α-tubulin (red) in male gametocytes at different time points after activation. DNA is stained with DAPI (blue); mpa = min post activation; scale bar = 5 µm. **(C)** Deconvoluted images improve the resolution of ARK2 and show its colocalization with spindle microtubules. Scale bar = 5 µm.

**Fig S3. The location of ARK2 and various subcellular markers**

**(A)** The location of ARK2-mCherry (red) and the kinetochore marker, NDC80-GFP (green) during male gametogony. DNA is stained with Hoechst dye (blue); scale bar = *5* μm. **(B)** Still images (at every 5 s) showing dynamic location of ARK2-mCherry and NDC80-GFP in gametocytes activated for 2 to 3 min. DNA is stained with Hoechst dye (blue); scale bar = 5 μm. **(C)** The location of ARK2-GFP (green) and the basal body and axoneme marker, kinesin-8B-mCherry (red) during male gametogony. DNA is stained with Hoechst dye (blue); scale bar = 5 μm. **(D)** Still images (at every 5 s) showing dynamic location of ARK2-GFP and kinesin-8B-mCherry in gametocytes activated for 4 to 5 min. DNA is stained with Hoechst dye (blue); scale bar = 5 μm.

**Fig S4. ARK2 associates with spindle microtubules**.

**(A)** STED confocal microscopy showing co-localization of ARK2 (purple) and α-tubulin (green) at spindle but not with cytoplasmic microtubules in gametocytes activated for 2 min. DNA is stained with SiR DNA (blue); scale bar = 1 μm. **(B)** Expansion microscopy showing co-localization of ARK2 (yellow) and α/β tubulin (purple) staining at spindle but not at cytoplasmic microtubules at 2 mpa. Scale bar = 1 μm. **(C)** 3D-SIM image showing locations of ARK2 (purple) and NDC80 (green) at 2 mpa. Scale bar = 1 μm. DNA (blue) is stained with DAPI.

**Fig S5. Generation and genotypic analysis of *Pb*ARK2-AID/HA and *P***_***clag-***_***ark2* parasites**.

**(A)** Schematic representation of auxin inducible degron (AID) strategy to generate *ARK2-AID/HA* parasites. (**B)**. Integration PCR of the ARK2-AID/HA construct in the *ark2* locus. Oligonucleotides used for PCR genotyping are indicated, and agarose gels to analyse the corresponding PCR products from genotyping reactions are shown. (**C)** ARK2-AID/HA protein expression level as measured by western blotting upon addition of auxin to mature purified gametocytes; α-tubulin served as a loading control. **(D)** Male gametogony (Exflagellation rate) of ARK2-AID/HA as measured upon addition of auxin and without auxin to mature purified gametocytes. **(E)** Schematic representation of the promoter swap strategy to construct *Pclag-ark2* parasites (placing ARK2 under the control of the clag promoter) by double homologous recombination. Arrows 1 and 2 indicate the primer positions used to confirm 5’ integration and arrows 3 and 4 indicate the primers used to confirm 3’ integration **(F)** Integration PCR of the promotor swap construct into the *ARK2* locus. Primer 1 (IntPTD245) and primer 2 (5’-IntPTD) were used to confirm successful integration of the selectable marker, resulting in a band of 460 bp. Primer 3 (3’-intPTclag) and primer 4 (IntPTD243) were used to determine the successful integration of the clag promoter, resulting in a band of 571 bp. Primer 1 (IntPTD245) and primer 4 (IntPTD243) were used to confirm a complete knock-in of the construct with a band at 4.5 kb and the absence of a band at 2.1 kb. **(G)** qRT-PCR showing normalised expression of ARK2 transcripts in *P*_*clag-*_*ark2* and WT-GFP parasites.

**Fig S6. Analysis of ookinete motility of *P***_***clag-***_***ark2* and WT-GFP parasites**

**(A)** Representative frames from time-lapse videos of WT-GFP and *Pclag-ark2* ookinetes in matrigel. Red arrow indicates the apical end of the ookinetes. Bar □= □5 µm. **(B)** Graph shows the quantitative data for WT-GFP and *Pclag-ark2* ookinete motility. (Error bar ± SD; n=3 independent experiments; >20 ookinetes were analysed for each experiment). **(C)** RNA sequence analysis showing downregulated transcript of ARK2 in *Pclag-ark2* parasites. **(D)** Gene ontology enrichment analysis showing the most affected genes involved in various biological processes.

**Fig S7. Generation of PbEB1-GFP parasites and analysis of PbEB1-GFP location during gametogony**

**(A)** Schematic representation of the endogenous *Pbeb1* locus, the GFP-tagging construct and the recombined *eb1* locus following single homologous recombination. Arrows 1 and 2 indicate the position of PCR primers used to confirm successful integration of the construct. **(B)** Diagnostic PCR of *eb1* and WT parasites using primers IntT264 (Arrow 1) and ol492 (Arrow 2). Integration of the EB1 tagging construct gives a band of 1267 bp. Tag = EB1-GFP parasite line. **(C)** Still images (at every 5 s) showing dynamic location of EB1-GFP in activated gametocytes at 1-2 min during male gametogony. DNA is stained with Hoechst dye (blue); scale bar = 5 μm. **(D)** Still images (at every 5 s) showing dynamic location of EB1-GFP in activated gametocytes at 2 to 3 mpa. DNA is stained with Hoechst dye (blue); Scale bar = 5 μm. **(E)** Still images (at every 5 s) showing dynamic location of EB1-GFP in activated gametocytes at 4 to 6 mpa. DNA is stained with Hoechst dye (blue); scale bar = 5 μm. **(F)** The location of EB1-GFP (green) and the kinetochore marker, NDC80-mCherry (red) during male gametogony. DNA is stained with Hoechst dye (blue); scale bar = *5* μm.

**Fig S8. EB1 associates with spindle microtubules**.

(A) 3D-SIM image showing location of EB1 (green) with NDC80 (purple) in gametocyte activated for 2 min and EB1 (green) with ARK2 (purple) in gametocytes activated for 4 min. 3D-SIM images showing location of EB1 (purple) and cytoplasmic SAS4 (green) in gametocyte activated for 2 min. DNA is stained with Hoechst dye (blue); scale bar = 1 μm. (B) STED confocal microscopy showing co-localization of EB1 (purple) and α-tubulin (green) at spindle but not with cytoplasmic microtubules in gametocytes activated for 2 min. DNA is stained with SiR DNA (blue); scale bar = 1 μm.

**Fig S9. PbEB1-GFP is located at the apical end of the parasite and at the putative MTOC and spindle like PbARK2-GFP during ookinete development** Live-cell imaging shows that EB1-GFP is located at the microtubule organising centre (MTOC) and spindles in the nucleus during ookinete development and then disappears in mature ookinetes (24 h). It is also located at the apical end of the growing protuberance during zygote to ookinete transition. A cy3-conjugated antibody, 13.1, which recognises the protein P28 on the surface of zygotes and ookinetes was used to mark these stages (red). Scale bar = 5 μm.

**Fig S10. Generation and genotypic analysis of** Δ***eb1* parasites**

**(A)** Schematic representation of the endogenous *eb1* locus, the targeting knockout construct and the recombined *eb1* locus following double homologous crossover recombination. Arrows 1 and 2 indicate PCR primers used to confirm successful integration in the *eb1* locus following recombination, and arrows 3 and 4 indicate PCR primers used to show deletion of the *eb1* gene. (**B)** Integration PCR of the *eb1* locus in WTGFP (WT) and knockout (Mut) parasites using primers: integration primer and ol248. Integration of the targeting construct gives band of expected size for each gene. **(C)** Gene ontology enrichment of upregulated genes in global transcriptomic analysis of Δ*eb1* gametocytes activated for 30 min, showing where the most affected genes are involved in various biological processes.

## Supplementary tables

**Table S1**. Overview of genomes and sequences used for generating Figure 1B.

**Table S2**. List of genes differentially expressed between *P*_*clag-*_*ark2* and WT-GFP gametocytes activated for 30 min.

**Table S3**. Spreadsheet (excel) file with unique peptide values for GFP-trap immunoprecipitate for gametocytes 1 minute after activation for WT-GFP, ARK2-GFP and EB1-GFP parasites. NAs are set to zero (0). Specific protein groups that belong to a similar functional class (e.g. replication machinery, kinetochore etc) are colour coded according to the scheme visualised in Fig 5B and Fig 8. Five parts of the table are present: (1) gene details I; containing gene name, manual annotations, amino acid number (AA) and molecular weight (MW), (2) correlations; Pearson (p) and Spearman (s, rank) correlation values for ARK2 and EB1, (3) PCA, principal components, (4) unique peptide values; NA is -, and * indicates that single peptide calls are to be approached with suspicion (minimal of 2 is usual cut-off), (5) gene details II; for GO terms and OG definitions that can be found at PlasmoDB (https://plasmodb.org/).

**Table S4**. List of genes differentially expressed between Δ*eb1* and WT-GFP gametocytes activated for 30 min

**Table S5**. Oligonucleotides used in this study.

## Supplementary Movies

**Video S1**. Time lapse video showing ARK2-GFP focal point extending to form a bridge-like spindle and breaking into two halves in gametocytes 1 to 2 min after activation. Still images used in Fig 2B.

**Video S2**. Time lapse video showing two ARK2-GFP bridge-like spindles breaking and producing four focal points in gametocytes 3 to 4 min after activation. Still images used in Fig S1G.

**Video S3**. Time lapse video showing four ARK2-GFP bridge-like spindles breaking and producing eight focal points in gametocytes 6 to 8 min after activation. Still images used in Fig S1H.

**Video S4**. Time lapse video showing ARK2-mCherry and NDC80-GFP dynamics in gametocytes activated for 1 to 2 min. Still images used in Fig 2D.

**Video S5**. Time lapse video showing ARK2-mCherry and NDC80-GFP dynamics in gametocytes activated for 2 to 3 min. Still images used in Fig S2D.

**Video S6**. Time lapse video showing ARK2-GFP and kinesin-8B-mCherry dynamics in activated gametocytes for 2-3 min. Still images used in Fig 2F.

**Video S7**. Time lapse video showing ARK2-GFP and kinesin-8B-mCherry dynamics in gametocytes activated for 4 to 6 min. Still images used in Fig S2F.

**Video S8**. Gliding motility of *Pclag-ark2* ookinetes. Still images used in Fig S4A

**Video S9**. Gliding motility of *WT-GFP* ookinetes. Still images used in Fig S4A

**Video S10**. Time lapse video showing EB1-GFP focal point extending to form a bridge like spindle in activated gametocytes for 1-2 min. Still images used in Fig S5C.

**Video S11**. Time lapse video showing EB1-GFP bridge breaking into two halves and accumulating at two focal points in a gametocyte activated for 2 to 3 min. Still images used in Fig S5D.

**Video S12**. Time lapse video showing two bridges of EB1-GFP breaking into four halves and accumulating at four focal points in a gametocyte activated for 2 to 3 min. Still images used in Fig S5E.

