## Supplementary figures and images for "*Plasmodium* ARK2-EB1 axis drives the unconventional spindle dynamics, scaffold formation and chromosome segregation of sexual transmission stages"

### Fig S1

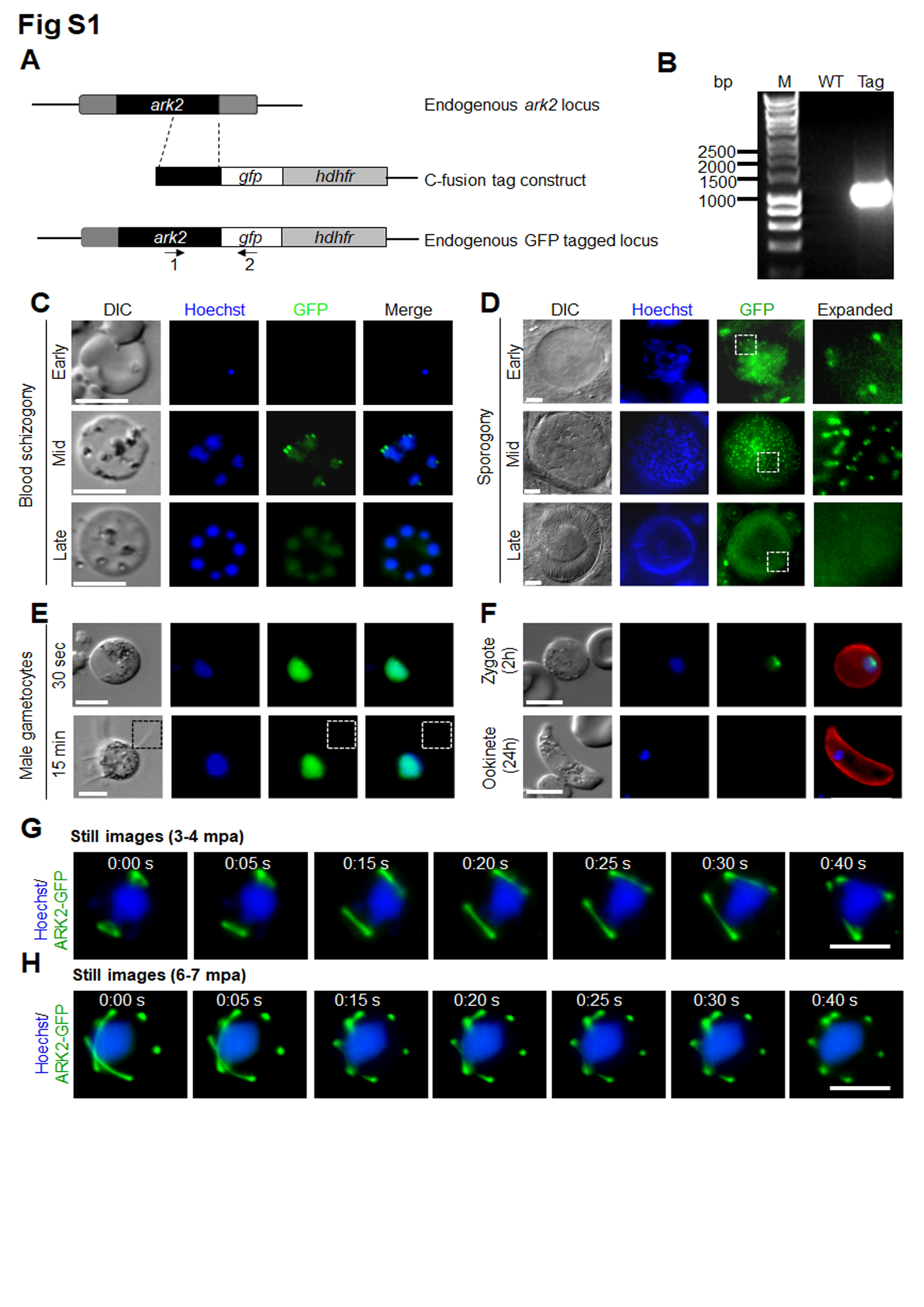

### Fig S2

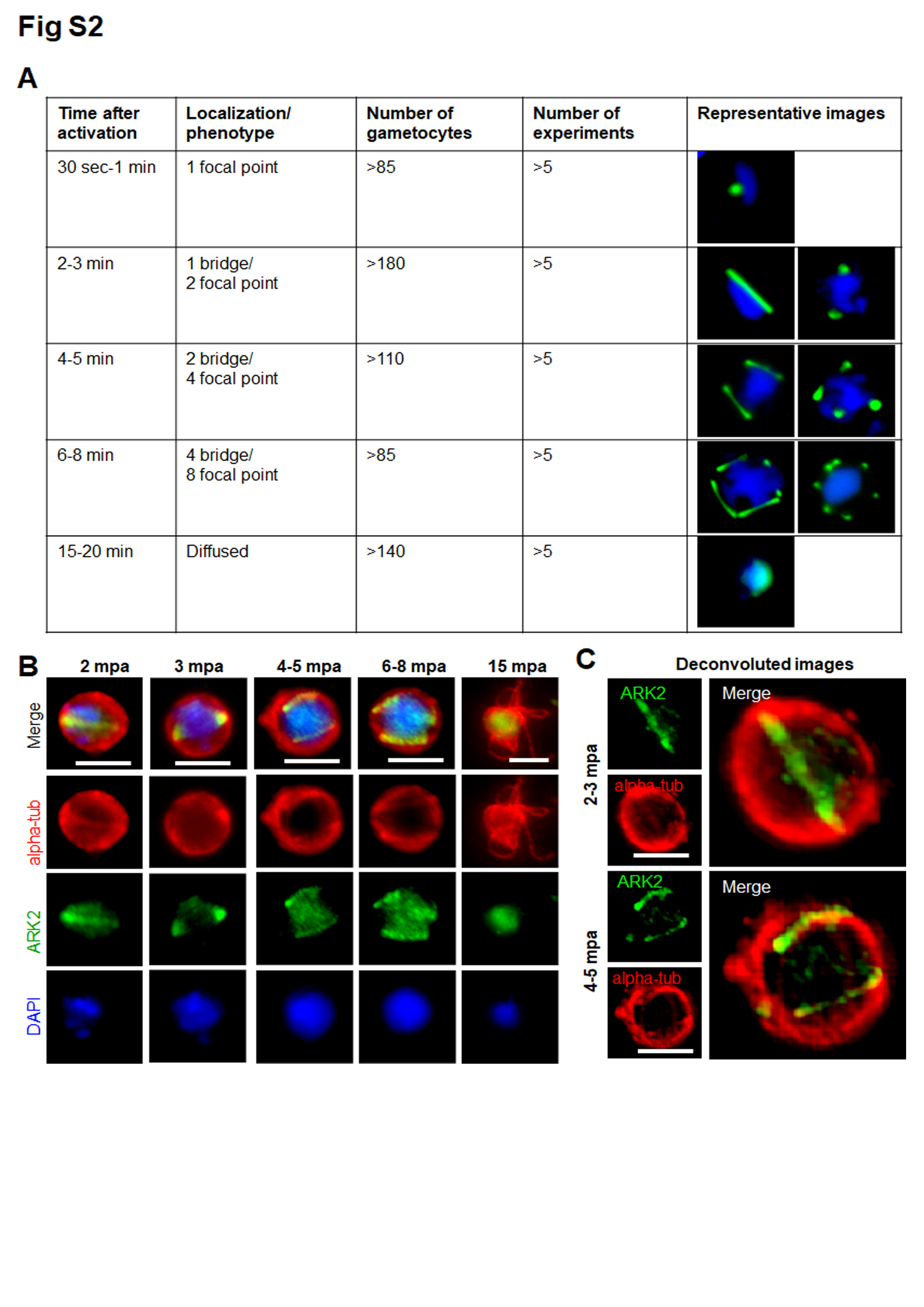

### Fig S3

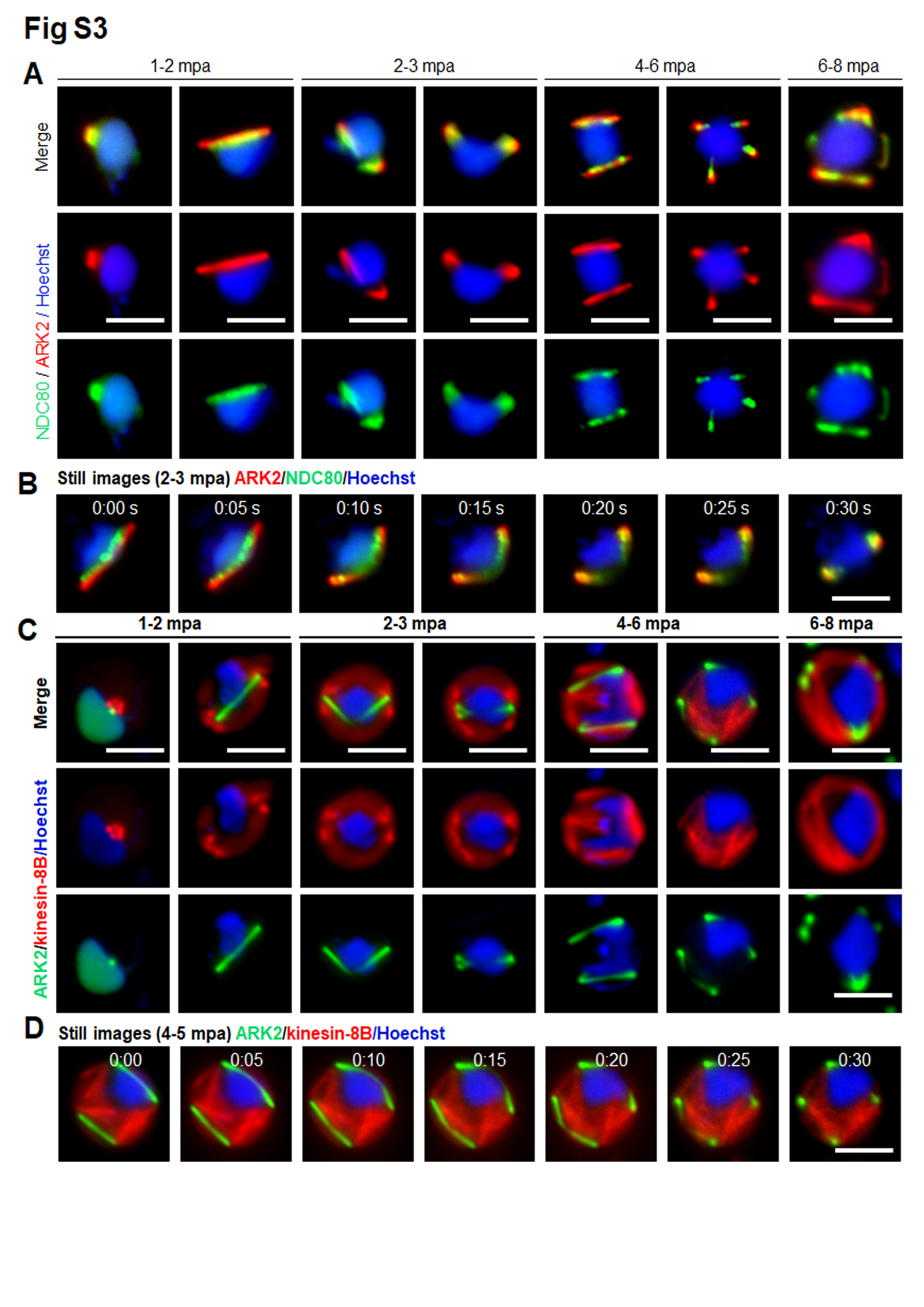

### Fig S4

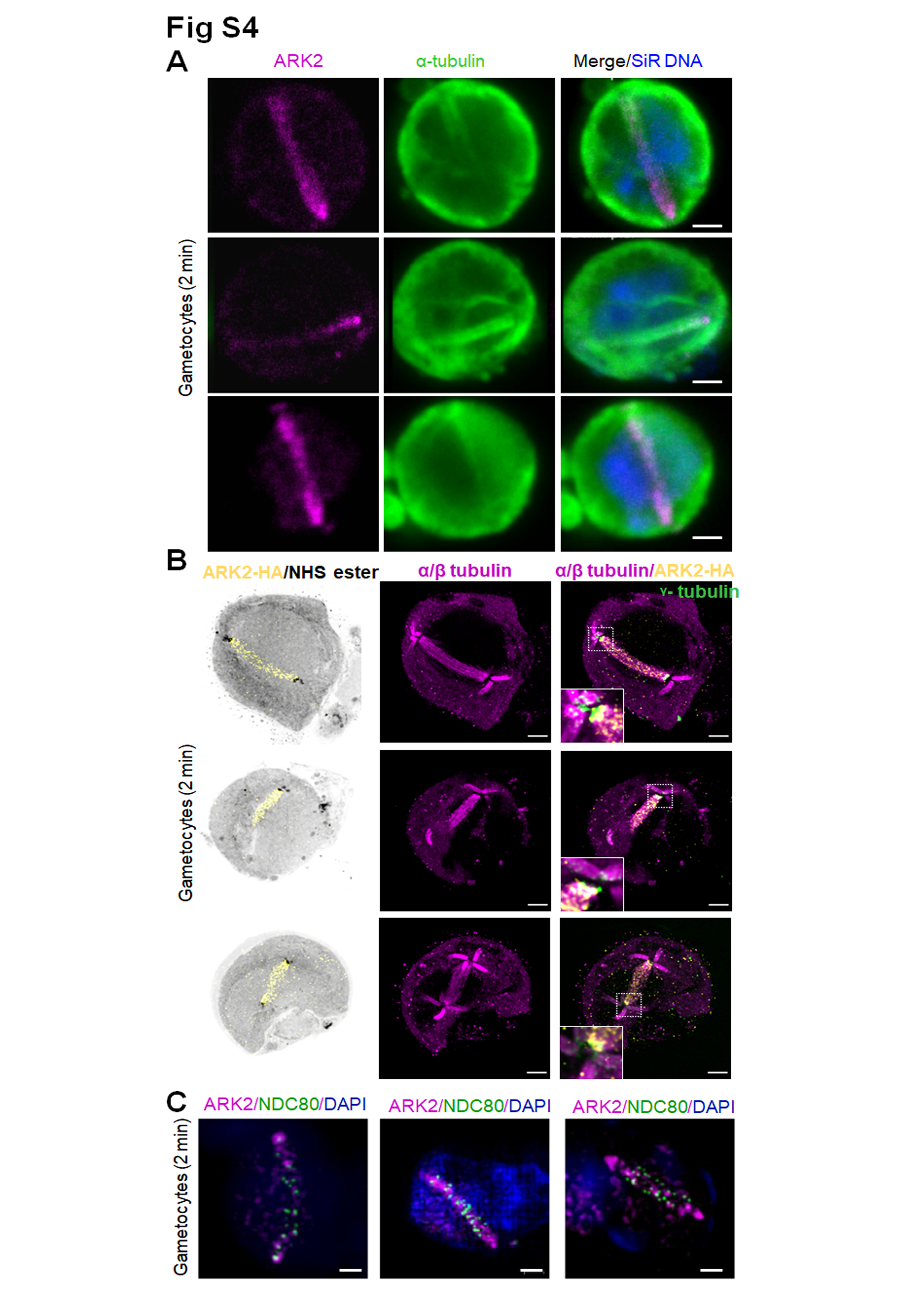

### Fig S5

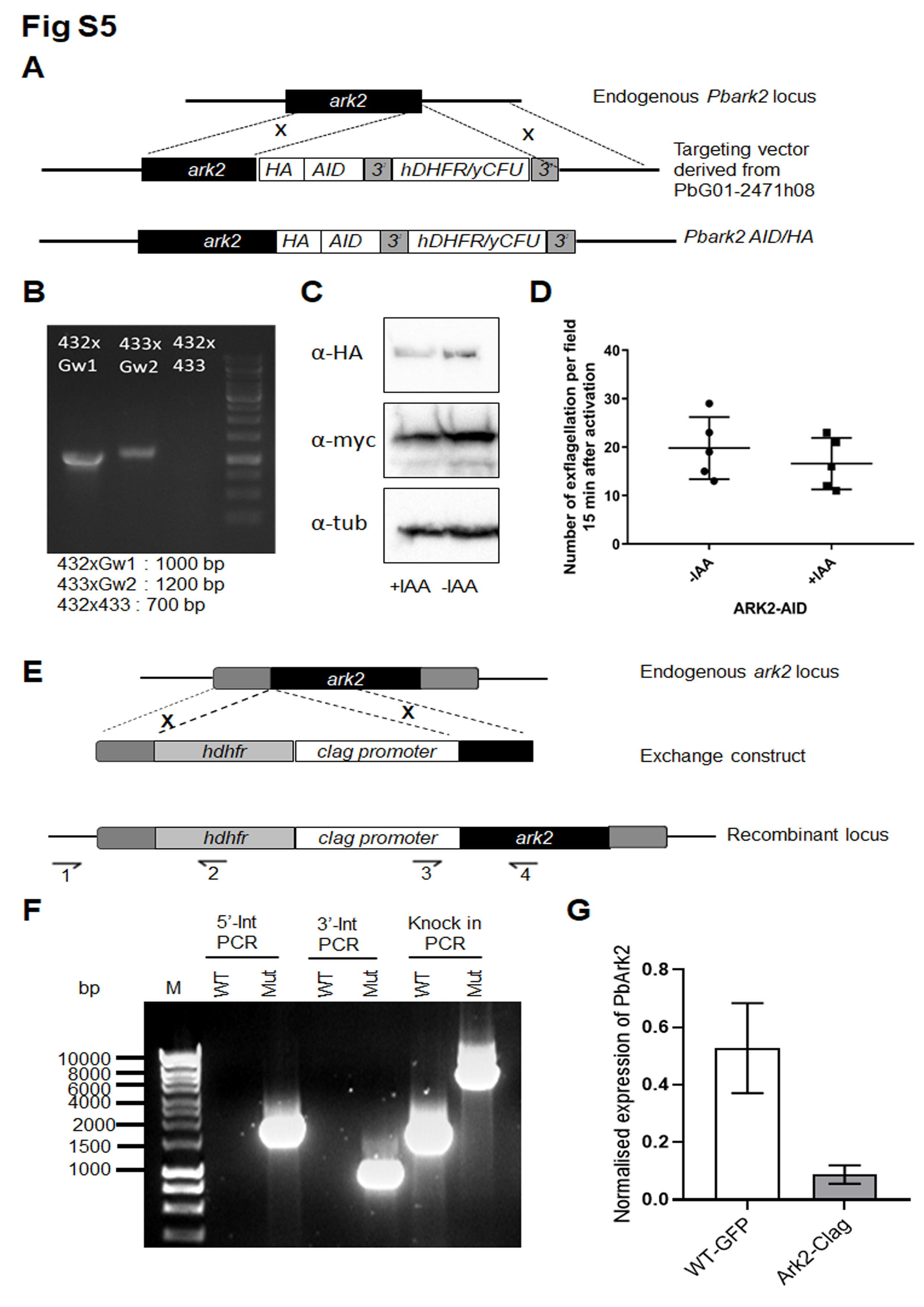

### Fig S6

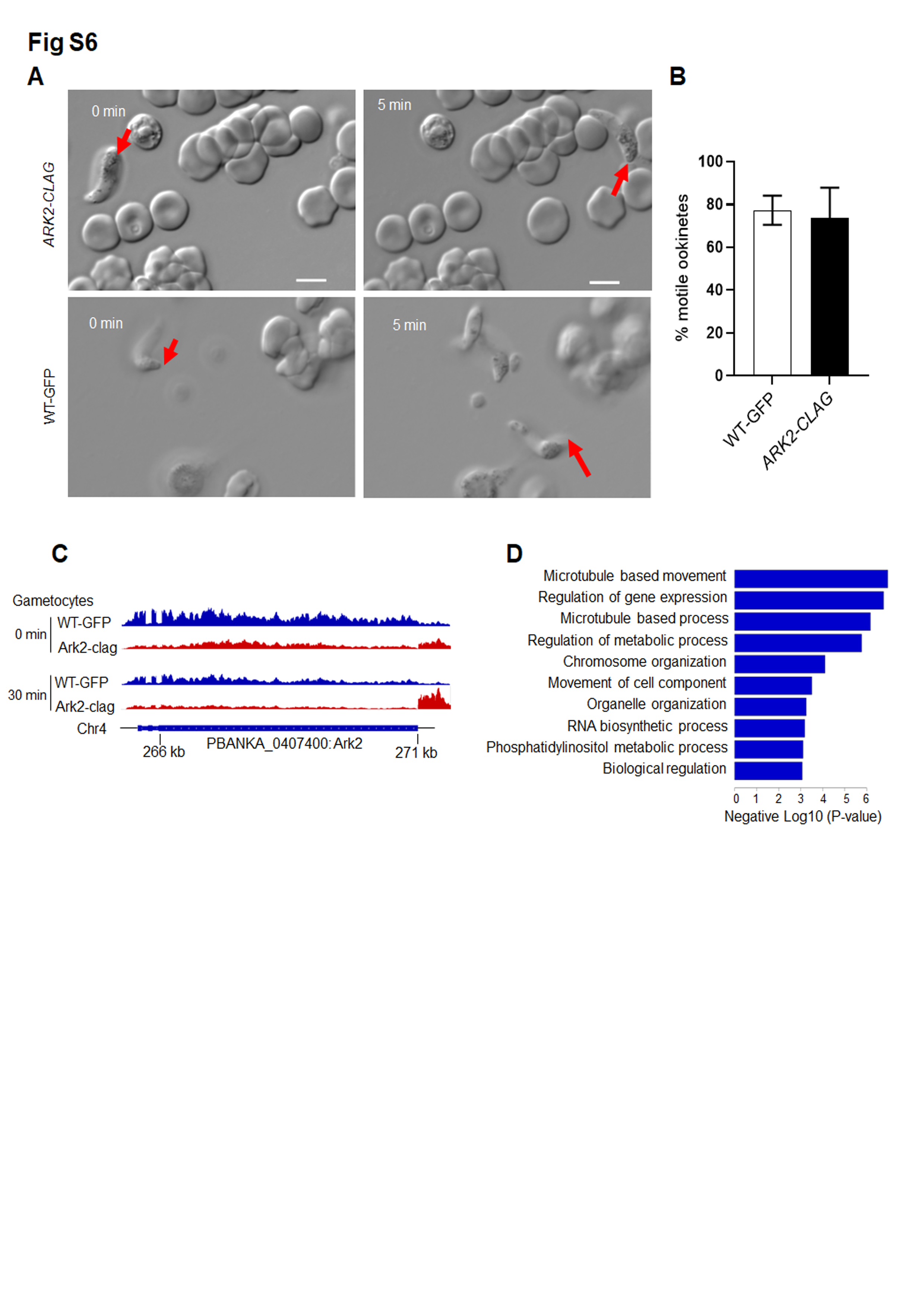

### Fig S7

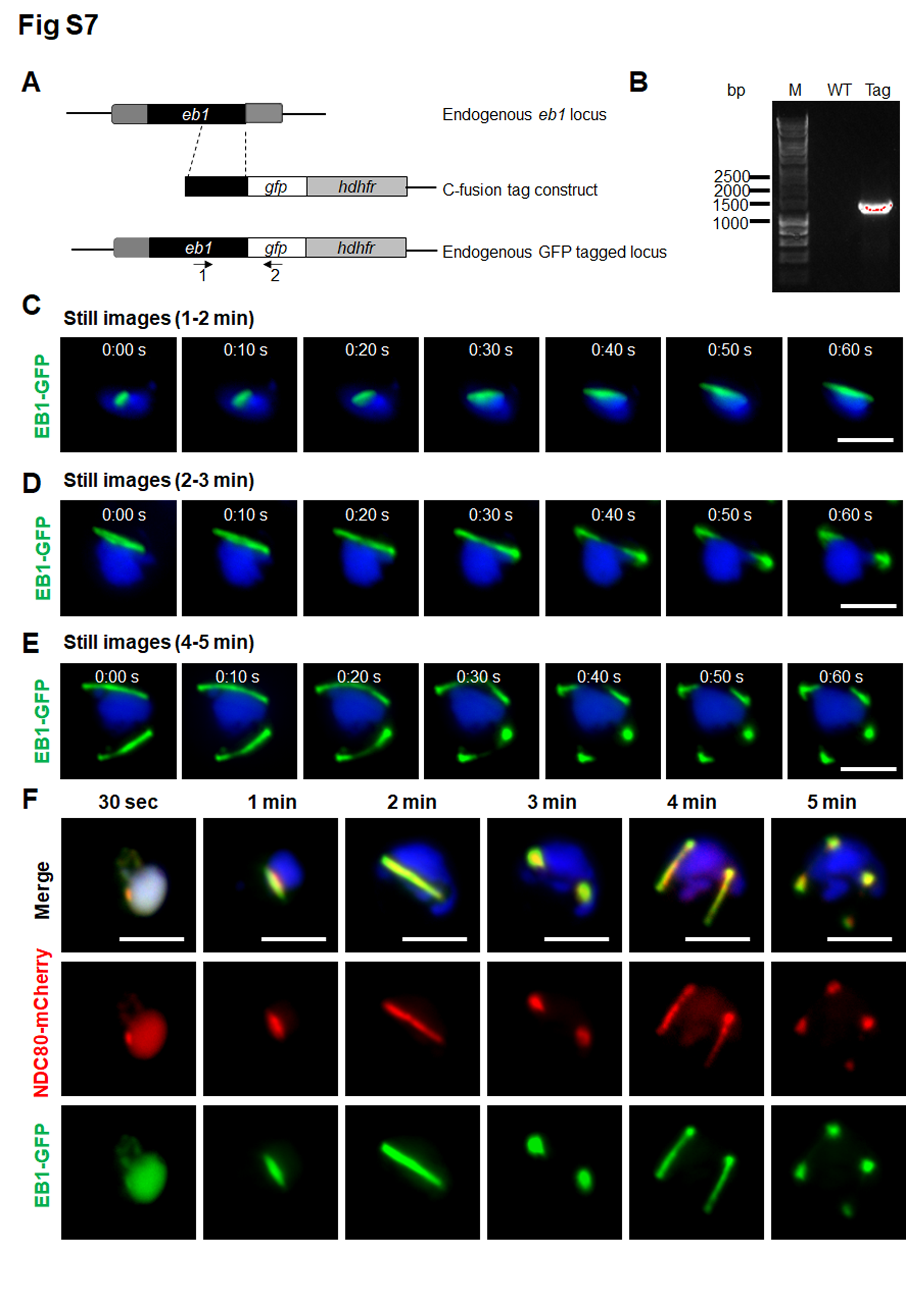

### Fig S8

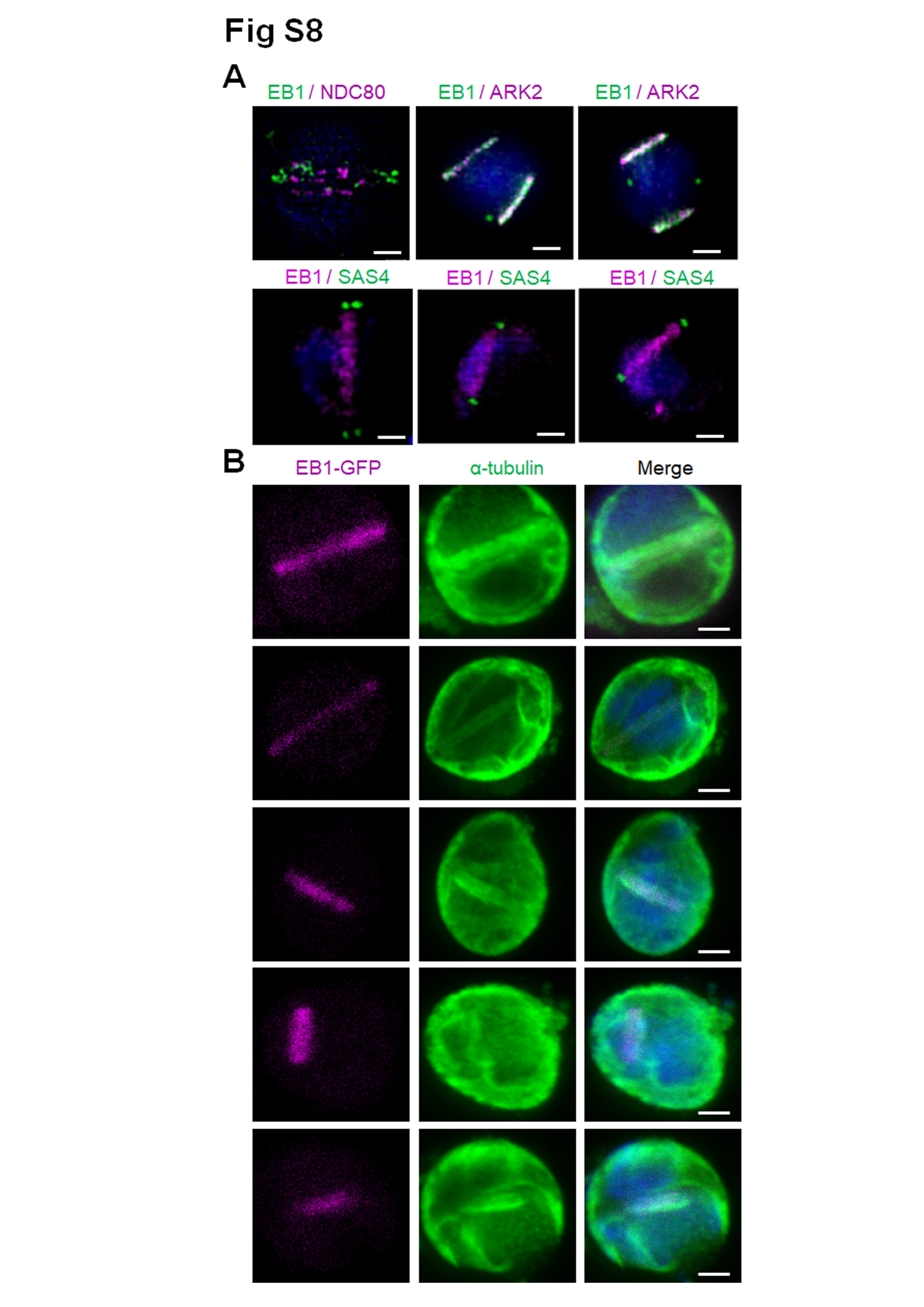

### Fig S9

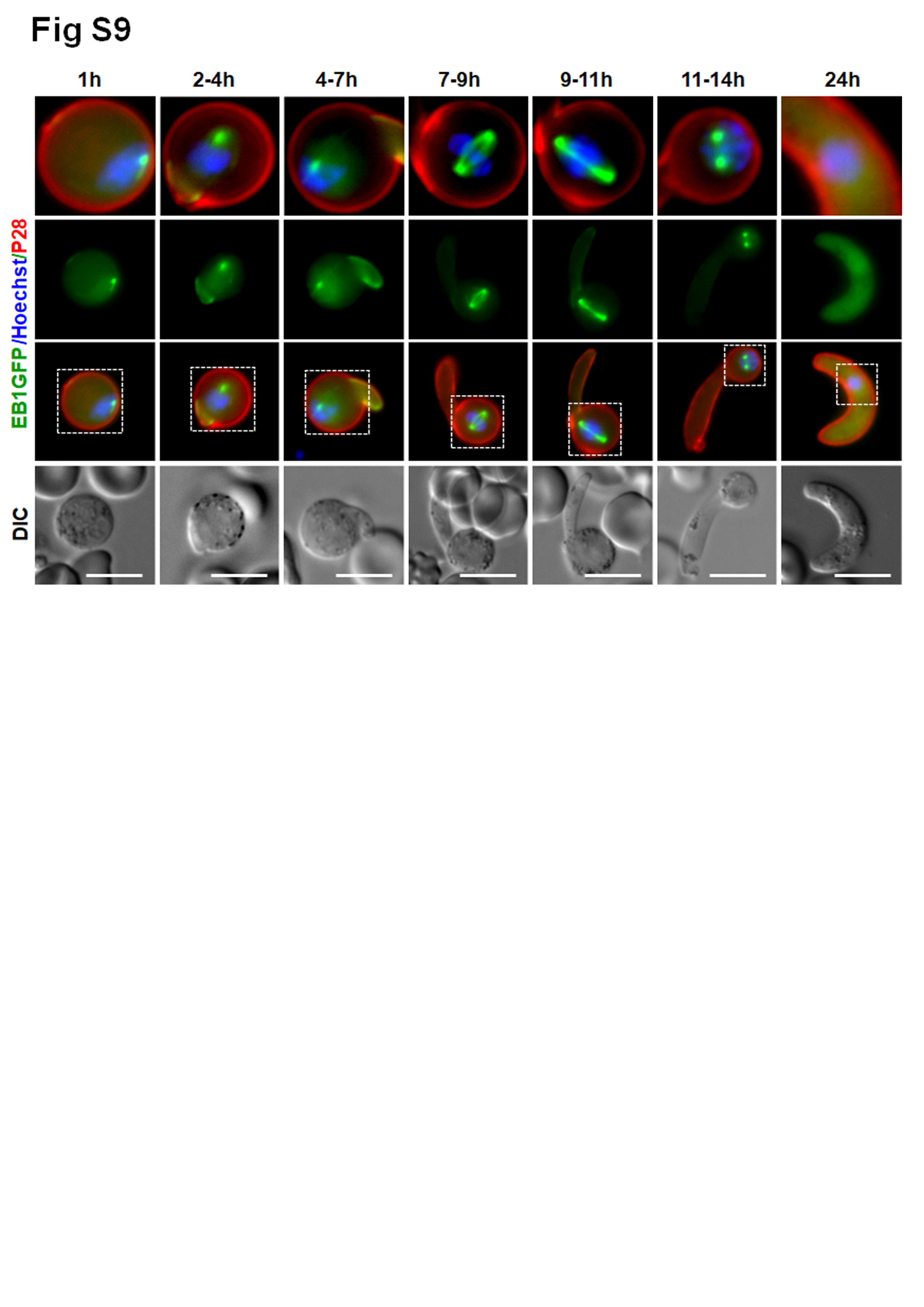
